# Dopamine-iron homeostasis interaction rescues mitochondrial fitness in Parkinson’s disease

**DOI:** 10.1101/2023.12.19.572356

**Authors:** Chiara Buoso, Markus Seifert, Martin Lang, Corey M. Griffith, Begoña Talavera Andújar, Maria Paulina Castelo Rueda, Christine Fischer, Carolina Doerrier, Heribert Talasz, Alessandra Zanon, Peter P. Pramstaller, Emma L. Schymanski, Irene Pichler, Guenter Weiss

**Author notes:** These authors have contributed equally (Irene Pichler, Guenter Weiss).

## Abstract

Imbalances of iron and dopamine metabolism along with mitochondrial dysfunction have been linked to the pathogenesis of Parkinson’s disease (PD). We have previously suggested a direct link between iron homeostasis and dopamine metabolism, as dopamine can increase cellular uptake of iron into macrophages thereby promoting oxidative stress responses. In this study, we investigated the interplay between iron, dopamine, and mitochondrial activity in neuroblastoma SH-SY5Y cells and human induced pluripotent stem cell (hiPSC)-derived dopaminergic neurons differentiated from a healthy control and a PD patient with a mutation in the α-synuclein (*SNCA*) gene. In SH-SY5Y cells, dopamine treatment affected the expression of transmembrane iron transporters and cellular iron accumulation. Furthermore, dopamine supplementation led to decreased mitochondrial respiration and reduced mitochondrial fitness, including reduced mtDNA copy number and citrate synthase activity, increased oxidative stress and impaired aconitase activity. In dopaminergic neurons derived from a healthy control individual, dopamine showed comparable effects as observed in SH-SY5Y cells. The hiPSC-derived PD neurons harboring an endogenous *SNCA* mutation demonstrated altered mitochondrial iron homeostasis, reduced mitochondrial capacity along with increased oxidative stress and alterations of tricarboxylic acid cycle linked metabolic pathways compared with control neurons. Importantly, dopamine treatment of these PD neurons promoted a rescue effect by increasing mitochondrial respiration, activating antioxidant stress response, and normalizing altered metabolite levels linked to mitochondrial function. These observations provide evidence that dopamine affects iron homeostasis, intracellular stress responses and mitochondrial function in healthy cells, while dopamine supplementation can restore this disturbed regulatory network in PD cells.

## Introduction

Parkinson’s disease (PD) is the second most common neurodegenerative disease after Alzheimer’s disease. PD is mostly an age-related disease, with incidence and prevalence increasing sharply with age (Collaborators, 2018; Pringsheim et al., 2014). The mean age of onset of PD is in the early-to-mid 60s, while early-onset PD refers to the minority of cases in which PD is diagnosed before 40 years of age (Bloem et al., 2021; Post et al., 2020). PD is characterized by dopaminergic neuronal loss in the *substantia nigra pars compacta* (*SNpc*), the formation of intracytoplasmic proteinaceous inclusions known as Lewy bodies (LBs), and iron accumulation (Forno, 1996; Jellinger et al., 1990; Riederer et al., 1989; Spillantini et al., 1998). Most PD cases are considered idiopathic (∼ 90%), and about 10% represent PD cases caused by mutations in single genes (Chang et al., 2017; Marras et al., 2016). Mutations in *SNCA, GBA1, LRRK2, PRKN, PINK1,* and *DJ-1*, among others, cause PD or early-onset forms of the disease (Blauwendraat et al., 2020). Point mutations and gene multiplications (duplication, triplication) in the *SNCA* gene encoding the α-synuclein protein (α-syn) cause severe forms of parkinsonism resulting in α-syn accumulation, oligomerization into fibrils, and LB formation (Chartier-Harlin et al., 2004; Shibasaki et al., 1995). α-syn accumulation is one of the molecular hallmarks of PD, together with mitochondrial dysfunction, oxidative stress and altered iron homeostasis (Bernal-Conde et al., 2019; Kaindlstorfer et al., 2018; Ma et al., 2021; Malpartida et al., 2021). Substantial evidence suggests that the mitochondrial electron transfer system, particularly Complex I (NADH:ubiquinone oxidoreductase) activity, is impaired in PD (Grunewald et al., 2019; Park et al., 2018). Moreover, mitochondrial DNA (mtDNA) alterations as well as mutations in genes involved in mitochondrial function (e.g. *PRKN*, *PINK1, SNCA*) can interfere with normal cell metabolism or cause rare genetic forms of the disease (Antonyova et al., 2020; Lang et al., 2022; Zanon et al., 2017). Specifically, a functional association between α-syn and mitochondrial membranes has been observed, with data suggesting that this interaction might induce α-syn accumulation and mitochondrial dysfunction (Choi et al., 2022; Monzio Compagnoni et al., 2020). Furthermore, oxidative damage takes part in the cascade of events leading to dopaminergic neurodegeneration in PD. Several sources can generate reactive oxygen species (ROS) within a cell, including interactions between redox-active metals, like iron, and oxygen species (Fenton and Haber-Weiss reactions) and activation of enzymes, such as nitric oxide synthase or NADPH oxidases (Dias et al., 2013; Koskenkorva-Frank et al., 2013), mediated by metals. Also, enzymatic oxidation or auto-oxidation of dopamine itself are recognized as causes of oxidative stress. Dopamine and its metabolites contain two hydroxyl residues that can induce cytotoxicity in dopaminergic neurons by producing highly reactive dopamine and DOPA quinones (Miyazaki and Asanuma, 2008). Outside of synaptic vesicles, dopamine is less stable and more prone to be metabolized via monoamine oxidase (MAO) or to auto-oxidize, resulting in the formation of cytotoxic ROS and neuromelanin (Sulzer et al., 2000), which can be protective or harmful depending on the amount of iron bound to it (Zucca et al., 2017). High levels of iron favor oxidative stress and therefore play a crucial role in the development of neurodegenerative diseases such as PD (Li et al., 2016; Ma et al., 2021).

Iron deposition was first observed in a PD brain by Lhermitte and colleagues in 1924 (Lhermitte et al., 1924), and it is still not clear whether this is a cause or a consequence of the disease process. However, iron accumulation in PD might be related to dopamine homeostasis (Hare and Double, 2016), and a disequilibrium between dopamine, iron and neuromelanin might trigger degeneration of dopaminergic neurons (Zucca et al., 2017). Furthermore, a correlation between iron overload in the brain, α-syn accumulation (Joppe et al., 2019; Ortega et al., 2016), and mitochondrial dysfunction (Munoz et al., 2016; Parihar et al., 2008) has been established. Iron metabolism is tightly regulated at systemic and cellular levels (Gao et al., 2019). Under physiological conditions, iron is an essential co-factor for metabolic processes, mitochondrial respiration, oxygen transport, and DNA synthesis or repair (Cheng et al., 2022; Cronin et al., 2019; Muckenthaler et al., 2017). In mitochondria, iron is necessary for heme synthesis, iron-sulfur (Fe-S) cluster biogenesis and can be stored in mitochondrial ferritin (Lane et al., 2015; Lill and Freibert, 2020). In the brain, iron is required for mitochondrial respiration and energy production, DNA replication, myelin synthesis, and as a co-factor of iron-containing enzymes like tyrosine hydroxylase and phenylalanine hydroxylase for neurotransmitter synthesis (dopamine, norepinephrine, and serotonin) (Cheng et al., 2022). Changes in iron status are known to regulate the dopaminergic system leading to or aggravating neurodegeneration (Ci et al., 2020; Matak et al., 2016). Because neuronal mitochondrial respiration is responsible for about 20% of the body’s oxygen consumption, the maintenance of physiological iron levels is critical for proper brain activity. Indeed, under physiological conditions, iron and dopamine collaborate for normal brain functioning, while under pathological conditions they can constitute a toxic couple (Hare and Double, 2016).

We have previously suggested a direct link between cellular iron homeostasis and dopamine metabolism, as dopamine can increase cellular uptake of iron into macrophages *in vitro* and *in vivo,* also promoting oxidative stress responses (Dichtl et al., 2019; Dichtl et al., 2018). Further, we have demonstrated that in subjects with restless legs syndrome, a neurological disease with indication of mitochondrial iron deficiency responding to dopaminergic medication like Levodopa, treatment with Levodopa resulted in improved mitochondrial function (Haschka et al., 2019). Based on these observations, we set out to further clarify the interaction of dopamine and iron in the context of PD by investigating the effect of dopamine on iron homeostasis, mitochondrial function and ROS production in neuroblastoma SH-SY5Y cells and PD-specific human neuronal models. We provide evidence that dopamine affects cellular iron trafficking, intracellular stress responses and mitochondrial function in neuronal cells. This regulatory network is compromised in PD cells but can be restored upon dopamine supplementation.

## Materials and Methods

### Cell lines and culture conditions

Neuroblastoma SH-SY5Y cells (ATCC, CRL-2266™) were cultured in Dulbecco’s modified Eagle’s Medium/Nutrient Mixture F-12+GlutaMAX™ supplement (DMEM/F12) (Thermo Fisher Scientific, 10565018) supplemented with 10 % fetal bovine serum and 1 % penicillin-streptomycin (Thermo Fisher Scientific, 15070-063). Human induced pluripotent stem cells (hiPSCs) from a control individual (hiPSC-SFC084-03-02, CTRL) and a PD patient with a triplication mutation of the *SNCA* gene (hiPSC-ND34391, 3x*SNCA*) were used. hiPSC-SFC084-03-02 was established through the StemBANCC consortium (https://cells.ebisc.org/STBCi033-B/), while hiPSC-ND34391 was obtained from the Coriell Cell Repository. Both lines were shared with the Institute for Biomedicine (Eurac Research) under MTA for research purposes. The study was approved by the Ethics Committee of the South Tyrolean Health Care System (approval number 102/2014 dated 26/11/2014 with extension dated 19/02/2020).

hiPSCs were cultured in mTeSR1 culture medium (Stem Cell Technologies, 85850) plus mTeSR1 supplement (Stem Cell Technologies, 85850) with 1 % Penicillin-Streptomycin (P/S) (Thermo Fisher Scientific, 15070-063) on Matrigel (Corning) coated plates under feeder-free conditions. hiPSC colonies were split every 3-4 days at a ratio of 1:5, the culture medium was changed every day. Cells were cultured at 37 °C, 5 % CO_2_ in a humidified atmosphere.

### Dopaminergic neuronal differentiation

The protocol for the differentiation was based on the floor-plate-based neural induction protocol reported by Kriks and colleagues (Kriks et al., 2011), with some minor variations as detailed in (Castelo Rueda et al., 2023; Zanon et al., 2017). Differentiations were started with a seeding density of 40,000 cells/cm^2^ for the 3x*SNCA* line and 50,000 cells/cm^2^ for the CTRL line. Neurons were collected at day 60 of differentiation to evaluate gene expression and protein levels or at day 35 for mitochondrial oxygen consumption assay.

### Treatment conditions for SH-SY5Y cells and dopaminergic neurons

Cells were treated with 5 µM ferric chloride (Sigma-Aldrich, 10025-77-1), 50 µM dopamine (Sigma-Aldrich, H8502), and dopamine in combination with ferric chloride at the same concentrations, 5 µM deferoxamine (DFO) (Sigma-Aldrich, 2604), or DFO in combination with dopamine, for 16 hours. To prevent endogenous dopamine degradation, 100 nM tranylcypromine (Calbiochem, 3980) was added to all treatment conditions.

### RNA extraction and quantitative real-time PCR (qRT-PCR)

Cells were lysed with TRIzol® Reagent (Thermo Fisher Scientific, 15596026), and RNA was extracted with the phenol-chloroform-based procedure or using Direct-zol RNA Miniprep Plus kit (Zymo research, R2070) following the manufacturer’s protocols. RNA was reverse transcribed using SuperScript VILO cDNA Synthesis Kit (Thermo Fisher Scientific, 11756050) and diluted to an RNA-corresponding concentration of 5 ng/μl. qRT-PCR was performed with 10 ng of cDNA on the CFX96 Real-Time PCR Detection System (Bio-Rad) either with the SsoAdvanced™ Universal Probes Supermix (Bio-Rad, 1725284) or SsoFast™ EvaGreen® Supermix (Bio-Rad, 1725200). Gene expression was calculated with the ΔΔCt method in the CFX96 Manager software (Bio-Rad).

### High-resolution respirometry (HRR) in neuroblastoma SH-SY5Y cells

High-resolution respirometry experiments were performed at 37 °C with continuous stirring (750 rpm) using the Oxygraph-2k (O2k, Oroboros Instruments, Innsbruck, Austria) as previously described (Castelo Rueda et al., 2023) with minor changes. After 16 hours of treatments, 1×10^6^ SH-SY5Y cells were resuspended in mitochondrial respiration medium MiR05 (Oroboros MiR05-Kit, Innsbruck, Austria). Oxygen consumption of permeabilized cells was measured applying the substrate-uncoupler-inhibitor titration protocol *SUIT-008 O2 ce-pce D025* (Lemieux et al., 2017). After measuring ROUTINE respiration, cells were permeabilized with optimum digitonin concentration (10 µg/ml (Doerrier et al., 2018)). Afterwards, NADH-linked substrates pyruvate (5 mM) and malate (2 mM) were added to evaluate the non-phosphorylating resting state LEAK state (PM_L_), followed by the addition of kinetically saturating concentration of ADP (2.5 mM) to assess OXPHOS capacity P (PM_P_). Cytochrome c (10 µM) was used to test the integrity of mitochondrial outer membrane (PMc_P_). Glutamate (10 mM) was injected as an additional substrate for the NADH-pathway (PGM_P_). Next, succinate was added to reconstitute convergent respiration through the NADH&succinate-pathway (PGMS_P_). Uncoupler titration (0.5 µM) of the protonophore carbonyl cyanide-p-trifluoromethoxy phenyl hydrazone, FCCP) was performed to obtain the electron transfer (ET) capacity E (noncoupled ET-state, PGMS*_E_*) and maximal respiration. Finally, residual oxygen consumption was evaluated after inhibiting Complex I and Complex III with rotenone (0.5 µM) and antimycin A (2.5 µM), respectively. We finally calculated the flux control ratio (FCR) by dividing the oxygen flow measurements acquired during the individual respiratory states by the one acquired in PGMS*_E_* state.

For living cells, 1×10^6^ SH-SY5Y cells cultured in DMEM/F12+GlutaMAX™ supplement were resuspended in mitochondrial respiration medium MiR05 (Oroboros MiR05-Kit, Innsbruck, Austria). Oxygen consumption of living cells was measured by applying the substrate-uncoupler-inhibitor titration protocol *SUIT-003 O2 ce-pce D020* (Doerrier et al., 2018). After measuring ROUTINE respiration, oligomycin (5 nM) was added to block the ATP synthase in order to reach the non-phosphorylating resting state (LEAK state, L(Omy)). Afterwards, we proceeded with uncoupler titrations of FCCP (0.1 μM steps) to obtain maximum electron transfer (ET) (noncoupled ET-state, E) measurements. Next, we blocked Complex I with Rotenone (0.5 μM) and evaluated the residual oxygen consumption (ROX) in the ROX state and the succinate pathway control state (S-pathway) upon succinate titration (10 mM). At the end, we assessed the ROX state by Antimycin A titration (2.5 μM). Respirometric measurements and data analysis were performed using the software DatLab 7.4 (Oroboros Instruments, Innsbruck, Austria). All reagents were purchased from Sigma-Aldrich. After the O2k run, cell suspensions were collected, snap-frozen, and stored at −80 °C to perform citrate synthase activity measurement.

### Oxygen consumption rate in hiPSC-derived neurons

Measurement of the oxygen consumption rate (OCR) in hiPSC-derived neurons was performed with the fluorescence-based Extracellular O_2_ Consumption Assay Kit (Abcam, ab197243) as described previously (Castelo Rueda et al., 2023). 200,000 cells/well were seeded on 96-well plates. Immediately after performing the OCR assay, the CyQuant™ proliferation assay (Thermo Fisher Scientific, C35012) was employed to determine the number of live cells in each well, according to the manufactureŕs instructions. OCR calculated for each well was normalized to the cell number determined by CyQuant fluorescence dye. OCR fluorescence intensities are expressed as relative fluorescence units (RFU) versus time (min).

### Mitochondrial DNA copy number

Genomic DNA was extracted with QIAamp DNA Blood Mini Kit (Qiagen, 51104) and quantified with NanoDrop 1000 (Thermo Fisher Scientific). 5 ng of DNA were used for evaluating the quantity of mitochondrial *MT-ND1* (VIC) and nuclear *B2M* (FAM) genes by qRT-PCR using TaqMan™ Gene Expression Assays (Thermo Fisher Scientific) with Taqman probes as described in (Wasner et al. 2022). Each sample was run in triplicate, and gene expression was calculated with the following formula: mtDNA copy number (CN)=2x[2^(ct^ ^B2M-ct^ ^ND1)^].

### Citrate synthase (CS) activity assay

Snap-frozen cell suspensions from O2k experiments were used for citrate synthase (CS) activity measurements, which were performed following the approach described in (Castelo Rueda et al., 2023). CS was assayed spectrophotometrically at 412 nm and 30 °C on an EnVision Multimode Plate Reader (Perkin-Elmer). All measurements were run in duplicate for each high resolution respirometry experiment, together with a blank control.

### Aconitase activity assay

Aconitase activity was measured with Aconitase Activity Assay Kit (Sigma-Aldrich, MAK051) according to manufacturer’s instructions. Briefly, 1.5×10^6^ cells were used for cell fractionation, and cytosolic as well as mitochondrial aconitase enzyme activities were measured as colorimetric signal in a coupled enzyme reaction. Absorbance was measured at 450 nm with the EnVision Multimode Plate Reader (Perkin-Elmer).

### Flow cytometry

Cells were stained for 15 min with 2.5 µM MitoSOX Red (Thermo Fisher Scientific, M36008) or for 30 min with 5 µM CellROX™ Green Flow Cytometry Assay Solution (Thermo Fisher Scientific, C10492) in fresh DMEM/F12+Glutamax w/o FBS and P/S. Cells were resuspended in 150 µl of FACS buffer (PBS supplemented with 0.5 % fetal bovine serum and 2 mM EDTA) containing DAPI (1:50,000, Sigma-Aldrich) as previously described (Fischer et al. 2021). Data were acquired using a CytoFLEX S flow cytometer (Beckman Coulter) and analyzed with FlowJo software (version 10.6.1, FlowJo LLC).

### Protein extraction and Western blotting

Western blot was performed as described previously (Castelo Rueda et al., 2023; Fischer et al., 2021). The following antibodies were diluted in 1 % non-fat dry milk in TBS-T: mouse anti-TFR1 antibody (1:1,000, Thermo Fisher Scientific, 13-6800, RRID:AB_2533029), rabbit anti-FPN antibody (1:2,000, Eurogentec, NRU 451443), rabbit anti-FTH (1:500, Sigma-Aldrich, F5012, RRID:AB_259622); rabbit anti-ACO2 (1:1,000, Abcam, ab129069, RRID:AB_11144142); rabbit anti-MFRN2 antibody (1:1,000, Bioss antibodies, bs-7157R), rabbit anti-FTMT antibody (1:1,000, Abcam, ab93428, RRID:AB_1056471), mouse anti-Nitrotyrosine antibody (1:1,000, Abcam, ab7048, RRID:AB_305725), mouse anti-GAPDH (1:10,000, Millipore, MAB374, RRID:AB_2107445), rabbit anti-PGC-1α (H300) (1:1,000, Santa Cruz Biotechnology, sc-13067, RRID:AB_2166218), rabbit anti-SOD2 (1:1,000, Cell Signaling Technology, 13141, RRID:AB_2636921), rabbit anti-β-actin (1:1,000; Sigma-Aldrich, A2066, RRID:AB_476693). Appropriate horseradish peroxidase (HRP)-conjugated secondary antibodies (1:2,000, Agilent Dako, P0448 polyclonal goat anti-rabbit immunoglobulins, RRID:AB_2617138; 1:4,000, Agilent Dako, P0447 polyclonal goat anti-mouse immunoglobulins, RRID:AB_2617137) were incubated in 1 % non-fat dry milk in TBS-T.

### Atomic absorption spectroscopy (AAS)

Intracellular iron content was measured by AAS as described previously (Theurl et al., 2016) using the Solaar M6 Dual Zeeman spectrometer (Thermo Scientific) with continuum source (QuadLine) and Zeeman background correction system.

### Extraction of intracellular metabolites

Metabolites were extracted as described previously (Gomez-Giro et al., 2019; Jager et al., 2016) with some minor modifications. Briefly, hiPSC-derived neuronal cell pellets were thawed on ice, and metabolites were extracted with water/methanol/chloroform, obtaining a three-phase system, where polar metabolites were enriched in the upper phase, non-polar metabolites in the lower phase, and cell debris constituted a solid interphase. The extraction fluid containing methanol (MeOH) and MilliQ H_2_O (4:1, v/v) was supplemented with seven internal standards (IS) (see Supplementary Material Section S1.1). Cells were homogenized with the extraction fluid, and then, the liquid-liquid extraction was performed by sequentially adding water/methanol/chloroform. The upper and lower phases were divided into separate tubes for the analysis of polar and non-polar metabolites and were evaporated to dryness using a Labconco CentriVap centrifugal vacuum concentrator at −4 °C overnight. Then, polar extracts were reconstituted in acetonitrile:MilliQ H_2_O (80:20, v/v), filtered with 0.2 μm PHENEX-RC 4 mm syringe filters, and transferred to Liquid Chromatography (LC)-vials, while non-polar extracts were reconstituted in toluene:methanol (1:9, v/v), centrifuged and transferred to LC-vials. Further details are given in the Supplementary Material Section S1 and Table S1.

### Liquid chromatography-high resolution mass spectrometry analysis

LC-high-resolution mass spectrometry (LC-HRMS) was carried out on a Thermo Scientific Vanquish Flex Quaternary LC coupled to a Thermo Q Exactive HF mass spectrometer. Two separate chromatographic runs were performed: hydrophilic interaction liquid chromatography (HILIC), for polar extracts, using the method reported in (Talavera Andujar et al., 2022) and reversed phase (RP), for non-polar extracts, adapting the method reported in (Cajka et al., 2017). Further details are given in the Supplementary Material, Section S1.2.

### Metabolomics data processing and analysis

Data processing was conducted using MS-DIAL (version 4.9.221218) (Tsugawa et al., 2020) as previously described (Talavera Andujar et al., 2022). For the analysis of polar fractions, the following libraries provided by MS-DIAL were used: MSMS_Public_EXP_Pos_VS17 and MSMS_Public_EXP_NEG_VS17 (16,481 unique compounds for MS/MS positive and 9,033 unique compounds for negative, respectively). Tentative identifications obtained from MS/MS spectral matching in MS-DIAL were compared against the retention times (RT), exact masses, and MS/MS of authenticated standards (**Table S1**) measured using the same method to obtain Level 1 confidence (i.e., confirmed structure) (Schymanski et al., 2014). Those features that were not present in the library were analyzed in MetFrag (Ruttkies et al., 2016) using the PubChemLite chemical database (31 March 2023 1.20.0 version) (Bolton et al., 2023; Schymanski et al., 2021) for tentative identification using *in silico* fragmentation.

For non-polar fractions, “lipidomics” was chosen as target omics in MS-DIAL, and an internal *in silico* database of MS-DIAL was used for tentative identification. Only features with MS/MS spectral matches with the MSP databases were considered for further analysis. Peaks were manually checked and additionally run in MetFrag for tentative identification using *in silico* fragmentation. Lipids satisfying these checks were classified with a confidence level 3a and were included for studying differences between the two cell lines and/or impact of iron and dopamine treatments. Criteria for confidence levels are reported in **Table S2**.

### Statistical analyses

Statistical analyses were performed with GraphPad Prism 9. When comparing treatments within one group, one-way ANOVA was used followed by Dunnett’s test to correct for multiple comparisons. For analyzing differences after treatments between groups in hiPSC-derived neurons, two-way ANOVA was used followed by Šídák’s test to correct for multiple comparisons. Mann Whitney test or a parametric unpaired t-test was used to compare differences between two groups. For the metabolite analyses, peak areas were normalized by sum of the total intensity (TIC). Statistical significance was tested using parametric unpaired t-test with Welch’s correction to assess metabolic differences between the control and the diseased cell lines (CTRL vs. 3x*SNCA*). For metabolites with different amounts between the two cell lines, a two-way ANOVA Šídák’s multiple comparisons test was performed to evaluate if iron or dopamine treatments have an impact on metabolite levels.

## Results

### Dopamine affects iron metabolism in SH-SY5Y cells

To investigate the effect of dopamine on cellular iron homeostasis in neuroblastoma SH-SY5Y cells, we examined the expression of critical cellular iron metabolism genes, including transferrin receptor 1 (*TFRC*, *TFR1*) and ferroportin (*SLC40A1*, *FPN*), which reflect the major avenues for cellular iron uptake and release, respectively (David et al., 2022; Muckenthaler et al., 2017), ferritin (FTH), the principal cellular iron storage protein consisting of heavy (FTH) and light chain subunits, as well as mitoferrin2 (*SLC25A28*, *MFRN2*), an iron transporter into mitochondria and good indicator of mitochondrial iron status (Ali et al., 2022; Muhlenhoff et al., 2015). Cells were treated with iron (FeCl_3_) or the iron chelator DFO for testing cellular responses to iron supplementation or depletion, and with dopamine alone or in combination with both compounds for analyzing their interaction. For iron and dopamine, different concentrations were tested. Based on our previous work (Dichtl et al., 2018), we screened for effects on *TFR1* mRNA expression and cellular toxicity after 16 hours of incubation. As a result, we selected 50 µM dopamine and 5 µM iron because higher iron concentrations resulted in cellular toxicity in this cell line, specifically when combined with dopamine (**Figure S1**). DFO showed anticipated effects on the regulation of iron genes, whereas iron, which was used at lower dosages as in other cellular systems and published reports (Dichtl et al., 2018), caused only small effects. Tranylcypromine was added to all treatments to prevent endogenous dopamine degradation, and an untreated condition was included to evaluate the basal expression of the investigated genes (Dichtl et al., 2019; Dichtl et al., 2018; Mercuri et al., 2000).

As expected, DFO treatment significantly increased *TFR1* mRNA expression (p=0.001) indicating effective chelation of cellular iron. However, DFO did not affect *FPN* mRNA expression. Interestingly, dopamine upregulated both *TFR1* (p=0.043) and *FPN* (p=0.011) mRNA levels (**Figure 1A**). Regarding mitochondrial iron status, dopamine significantly upregulated *MFRN2* (p=0.042) (**Figure 1A**). To exclude unspecific effects of dopamine, FeCl_3_, and DFO or combinations thereof, we analyzed cellular viability, which showed no significant differences between untreated and treated cells (**Figure S2**).

**Figure 1.**
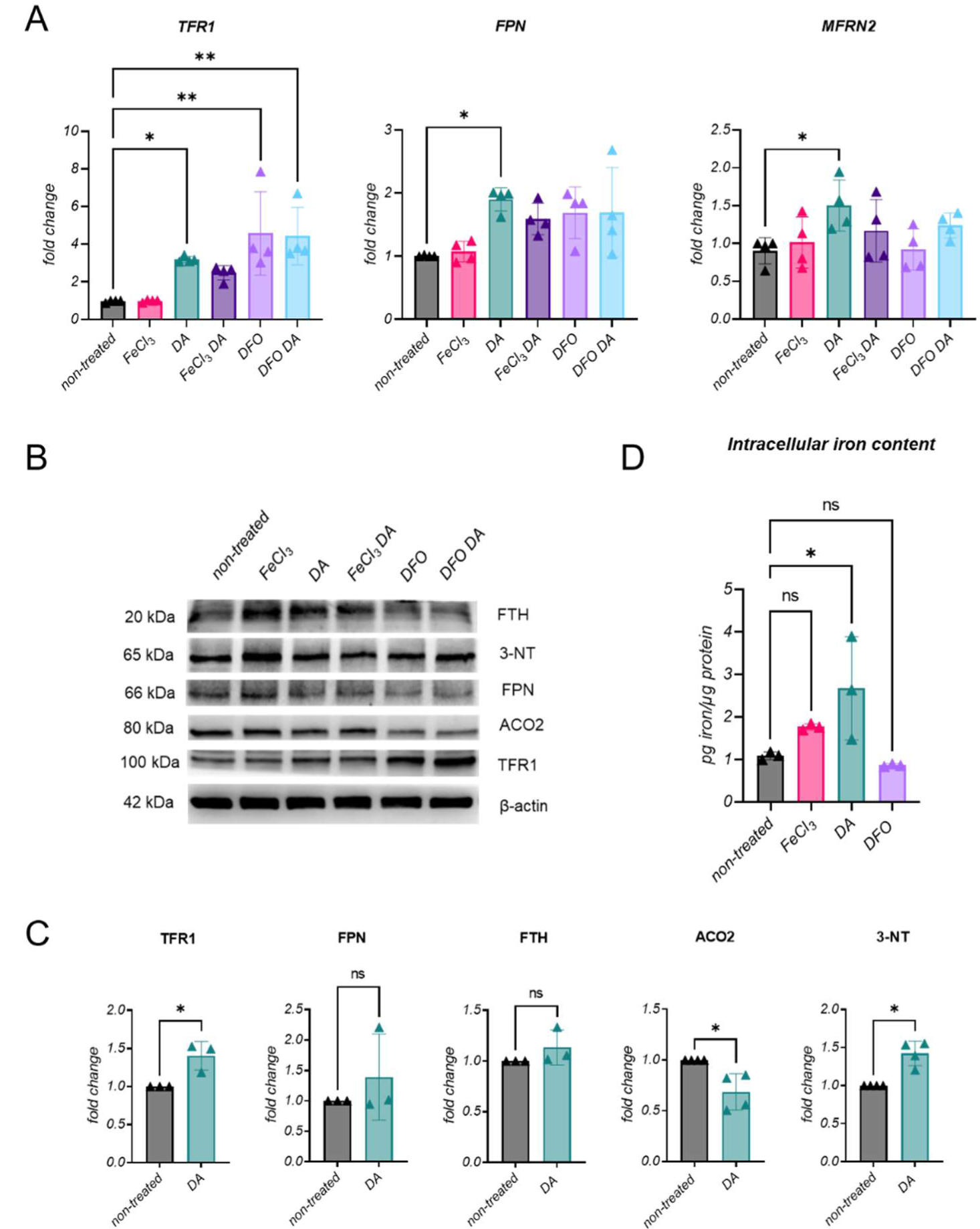
Expression levels of genes and proteins involved in iron metabolism and intracellular iron levels in SH-SY5Y cells. A) TFR1, FPN, MFRN2 mRNA expression levels normalized with TUBA1A (tubulin); n=4. B) Representative western blot showing TFR1, FPN, FTH, ACO2, 3-NT, protein levels upon treatments; β-actin was used as a loading control. Molecular mass markers are in kilodaltons (kDa). C) Densitometric analyses of TFR1, FPN, FTH, ACO2 and 3-NT protein levels after treatment with dopamine; n≥3. Statistical differences were calculated by Mann Whitney test. *p≤0.05. D) Intracellular iron levels measured by atomic absorption spectroscopy in cells treated with dopamine, iron and DFO. Results were normalized to protein content; n=3. Statistical differences were calculated by ordinary one-way ANOVA followed by Dunnett’s test to correct for multiple testing. *p≤0.05; **p≤0.01;****p≤0.0001.

To investigate whether the observed effects on iron-metabolism-related gene expression translated into corresponding changes of protein expression, we quantified protein levels by means of western blotting. Dopamine treatment resulted in increased TFR1 protein levels (p=0.029), in a similar way as DFO treatment (p=0.012) and their combination (p=0.032), further suggesting that dopamine supplementation may result in promotion of iron uptake possibly as a consequence of iron binding by dopamine (Dichtl et al., 2018) (**Figure 1B, C; Figure S3A**). Of note, while iron resulted in the anticipated increase of FTH expression, no changes were observed with dopamine neither on the mRNA nor the protein level. For FPN, no dopamine-induced upregulation on the protein level was observed (**Figure 1B, C; Figure S3B**).

In addition, we evaluated protein expression of mitochondrial aconitase (mitochondrial, m-aconitase, ACO2), an iron-sulfur (FeS) cluster containing enzyme that catalyzes the interconversion of citrate to iso-citrate as part of the tricarboxylic acid (TCA) cycle (Beinert and Kennedy, 1993), and because its translation is controlled by cellular iron availability (Muckenthaler et al., 2017). As an indicator of oxidative and nitrosative stress responses in the cell, we also determined nitrotyrosine (3-NT) (Kuhn et al., 2004). Protein levels of ACO2 were found to be significantly decreased upon dopamine (p= 0.029) treatment, which would be in line with translational repression by iron deficiency as described. Of note, dopamine treatment resulted in higher 3-NT levels, indicative for intracellular oxidative stress, which can also impair ACO2 expression (Liang and Patel, 2004) (**Figure 1B, C; Figure S3A**).

Since the data on gene and protein expression suggest alterations in cellular iron status after dopamine treatment, intracellular iron levels were measured by AAS. Dopamine treatment was associated with a significant increase of total intracellular iron levels (p=0.032). FeCl_3_ and DFO treatments resulted in increased and decreased intracellular iron measurements, respectively, as expected (**Figure 1D**).

### Dopamine affects mitochondrial fitness in SH-SY5Y cells

Since the data collected thus far indicated that dopamine treatment of SH-SY5Y cells altered cellular iron homeostasis, ACO2 protein levels and oxidative stress response, we questioned whether dopamine directly affects mitochondrial function. We measured mitochondrial respiration by HRR in living and permeabilized SH-SY5Y cells upon treatment with dopamine, FeCl_3_ and the combination of both compounds, because iron is necessary for mitochondrial function (Volani et al., 2017). Representative traces of real-time oxygen consumption illustrate the multiple substrate-uncoupler-inhibitor titration (SUIT) protocol in permeabilized SH-SY5Y cells (**Figure 2A**).

**Figure 2.**
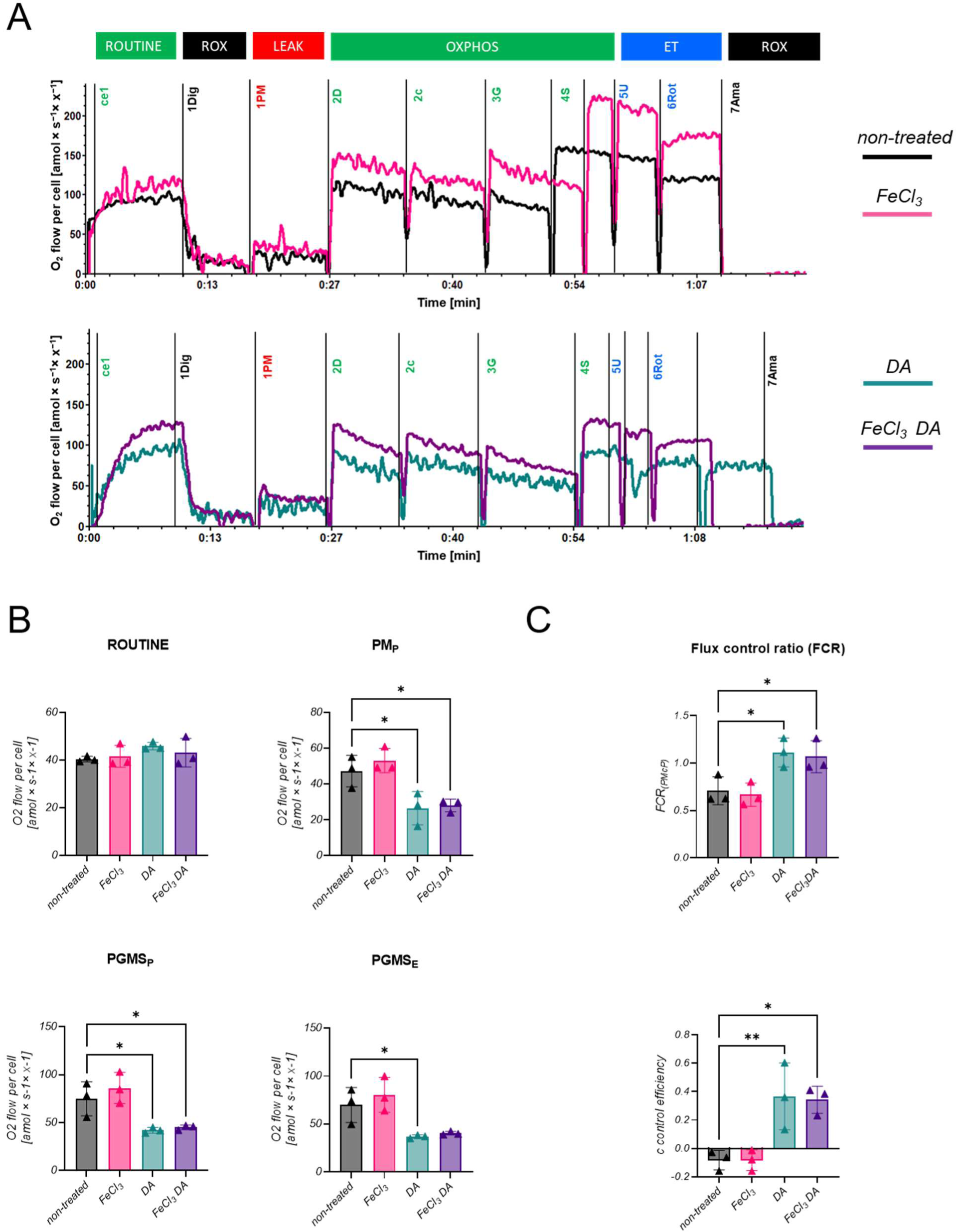
Effect of dopamine and FeCl_3_ on mitochondrial respiration in permeabilized SH-SY5Y neuroblastoma cells. A) Representative traces of oxygen consumption in permeabilized SH-SY5Y cells treated with the compounds of interest for 16 hours with sequential titrations. Respiration values were expressed as 0_2_ flow per cell [amol × s^−1^ × x^−1^]; y-axis) as a function of time (x-axis). Sequential steps: addition of cells (ce1), digitonin (1Dig, 10 µg/ml), pyruvate&malate (1PM, 5 mM&2 mM), ADP (2D, 2.5 mM), cytochrome c (2c, 10 µM), glutamate (3G, 10 mM), succinate (4S, 10 mM), FCCP (5U, 0. 5 µM step), rotenone (6Rot, 0.5 µM), antimycin A (7Ama, 2.5 μM). Non-treated: black; FeCl_3_: pink; DA: turquoise; FeCl_3_ and DA: purple. B) Oxygen consumption in permeabilized SH-SY5Y cells treated with the compounds of interest for 16 hours. ROUTINE: ROUTINE respiration in the physiological coupling state controlled by cellular energy demand, energy turnover and the degree of coupling to phosphorylation; PM_P_: OXPHOS capacity with PM as NADH-linked substrates; PGMS_P_: OXPHOS capacity of the convergent NADH&succinate-pathway into Q; PGMS_E_: noncoupled ET-state of the NADH&succinate-pathway. Each condition was measured in three independent assays. C) Flux control ratio after cytochrome c titration (FCR_(PMcP)_) and cytochrome c control efficiency calculated as 1-PM_p_/PMcP. Statistical analysis was carried out by using the ordinary one-way ANOVA followed by Dunnett’s test to correct for multiple testing. *p≤0.05, **p≤0.01.

In living (non-permeabilized) cells, differences upon dopamine treatment were evident in the maximal respiration (ET), indicating a dyscoupling (**Figure S4**). In permeabilized cells treated with dopamine or FeCl_3_ combined with dopamine, we observed a decreased oxygen consumption after titration of kinetically saturating concentrations of ADP in the presence of pyruvate and malate (PM_P_) (p=0.023 and p=0.033, respectively), and succinate (PGMS_P_) (p=0.033 and p=0.034, respectively). In addition, cells treated with dopamine alone showed a significant decrease in maximal oxygen flux (PGMS_E_) (p=0.036) (**Figure 2B**).

Given the observed decrease in OXPHOS and ET capacity, we calculated the flux control ratio (FCR) to investigate whether mitochondrial quality defects were induced by treatments. After cytochrome c titration, FCR was significantly increased by dopamine (p=0.026) and its combination with FeCl_3_ (p=0.045) (FCR_(PMcP)_). Additionally, the treatments increased also cytochrome c control efficiency (c control efficiency) suggesting an alteration of the integrity of the outer mitochondrial membrane (dopamine: p=0.010, dopamine and FeCl_3_: p=0.013) (**Figure 2C**). These observations suggest that dopamine changes mitochondrial quality, which may contribute to the observed decrease in mitochondrial respiration.

### Dopamine affects mitochondrial numbers and induces oxidative stress in SH-SY5Y cells

To further investigate mitochondrial homeostasis, we measured mitochondrial DNA (mtDNA) copy numbers in SH-SY5Y cells treated for 16 hours with FeCl_3_, dopamine, and the combination of FeCl_3_ and dopamine. We detected a significant drop of mtDNA copy number upon treatment with dopamine (p=0.022) and for concomitant dopamine with FeCl_3_ supplementation (p=0.010) (**Figure 3A**). Compared to untreated conditions, mtDNA copy number decreased by 17.97 % after FeCl_3_ treatment, by 38.56 % after dopamine treatment, and by 44.77 % after the treatment with iron and dopamine. To confirm these data, we measured citrate synthase activity, which is a functional marker of the amount of mitochondria, and found a decrease for all treatments compared to untreated cells, comparable to observations for the mtDNA copy number (FeCl_3_ p=0.011; DA p=0.002; FeCl_3_+DA: p=0.014). Together, these data indicate a reduced mitochondrial content upon treatment with dopamine, independent of FeCl_3_ supplementation (**Figure 3B**).

**Figure 3.**
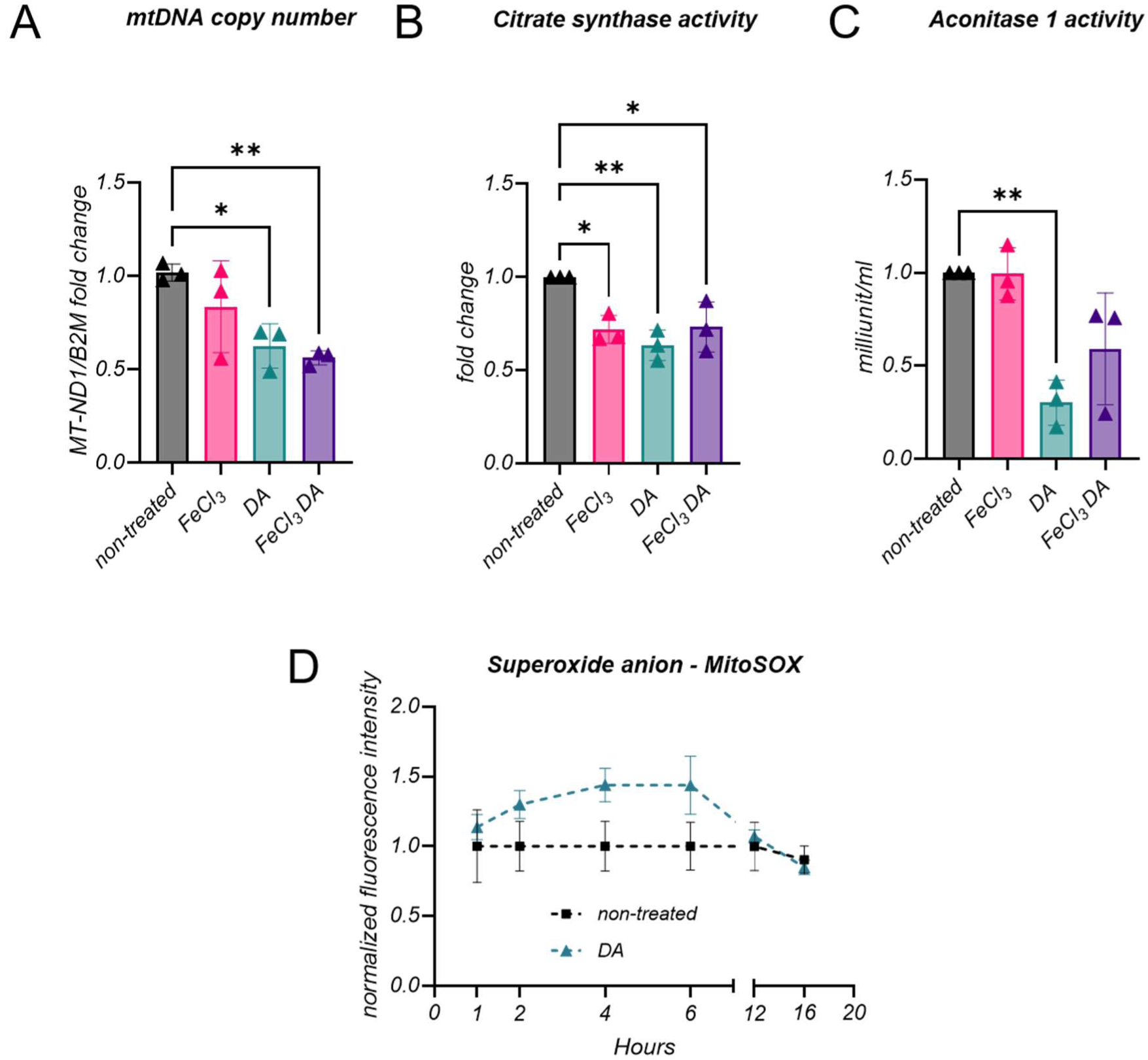
Evaluation of mitochondrial content and oxidative stress. A) mtDNA copy number measured by RT-PCR of MT-ND1, normalized to B2M gene on genomic DNA of permeabilized cells after 16 hours of treatment. B) Citrate synthase activity in permeabilized cells measured after 16 hours of treatment. C) Cytosolic (c-) aconitase activity upon treatment with the compounds of interest. D) MitoSOX Red signal was quantified at different time points in non-treated conditions and upon dopamine treatment. Successively, the signal was normalized for non-treated cells. At each time point, cells treated with dopamine are compared to cells without treatment as control. Each data point represents the mean of 3 measurements ± standard deviation (SD). Statistical analysis was carried out by using the ordinary one-way ANOVA followed by Dunnett’s test to correct for multiple testing. *p≤0.05; **p≤0.01.

Next, we measured aconitase activity to investigate whether the detrimental effect of dopamine on mitochondrial respiration and content could be mediated by oxidative stress. Aconitase 1 (cytosolic, c-aconitase, ACO1) and mitochondrial ACO2 are reversibly inactivated by oxidative stress and can therefore be used as a marker for oxidative damage (Rouault, 2006). Compared to the untreated condition, a significant decrease of ACO1 activity after dopamine treatment was detected (p=0.003), and the same trend was observed when combining dopamine with FeCl_3_ (p=0.053), while FeCl_3_ alone did not show an effect (**Figure 3C**). Although ACO2 protein expression was reduced (**Figure 1**), we only observed a non-significant reduction of its activity upon dopamine and the combined treatment (**Figure S5**).

To measure total cellular ROS and mitochondrial ROS (mROS) production, cells were stained with CellROX dye, which upon oxidation by ROS exhibits a photostable green fluorescence, and MitoSox Red dye, which specifically targets mitochondria in live cells and, upon oxidation by superoxide anion, emits a red fluorescence signal. While mitochondrial superoxide anion levels remained stable for up to 16 hours in untreated cells, they started to rise after 2 hours of dopamine treatment, reaching maximum levels at 4 to 6 hours, and then dropping to baseline levels. This suggests subsequent activation of antioxidant systems in the cells that protect remaining mitochondria from oxidative stress or the removal of damaged mitochondria (**Figure 3D**). A similar trend was detected for treatments with FeCl_3,_ dopamine and FeCl_3_, and with DFO (**Figure S6A**). Moreover, CellROX signals, measuring generalized oxidative stress, followed a very similar pattern upon the different treatments (**Figure S6B**). ROS might trigger mitochondrial damage responsible for the decrease in mitochondrial content and quality resulting in an overall reduction of mitochondrial respiration.

### Dopamine affects iron metabolism in hiPSC-derived neurons

We then studied whether the effects of dopamine on SH-SY5Y cells also apply to hiPSC-derived dopaminergic neurons of a control and a PD patient-derived cell line carrying a triplication mutation of the *SNCA* gene (3x*SNCA*). Neurons were treated for 16 hours with the compounds of interest (FeCl_3_, DFO, dopamine or their combination) and mRNA expression of *TFR1*, *FPN,* and *MFRN2* was evaluated. Comparison of baseline mRNA expression between control and 3x*SNCA* hiPSC-derived neurons showed that *TFR1* and *FPN* were significantly increased in the PD line (p=0.0002 and p=0.005, respectively), suggesting an altered iron status in the 3x*SNCA* line, while the mitochondrial marker *MFRN2* did not differ between control and PD lines (**Figure 4A**). After dopamine treatment, there was a trend for an increased mRNA expression of *TFR1* in the control line compared to non-treated cells, which was significant in the PD line (p=0.032). *FPN* levels showed minor differences among groups in both lines. Interestingly, for *MFRN2* there were no differences upon treatments in the control line, while in the patient line, dopamine treatment significantly reduced its mRNA expression (p=0.011) (**Figure 4B**).

**Figure 4.**
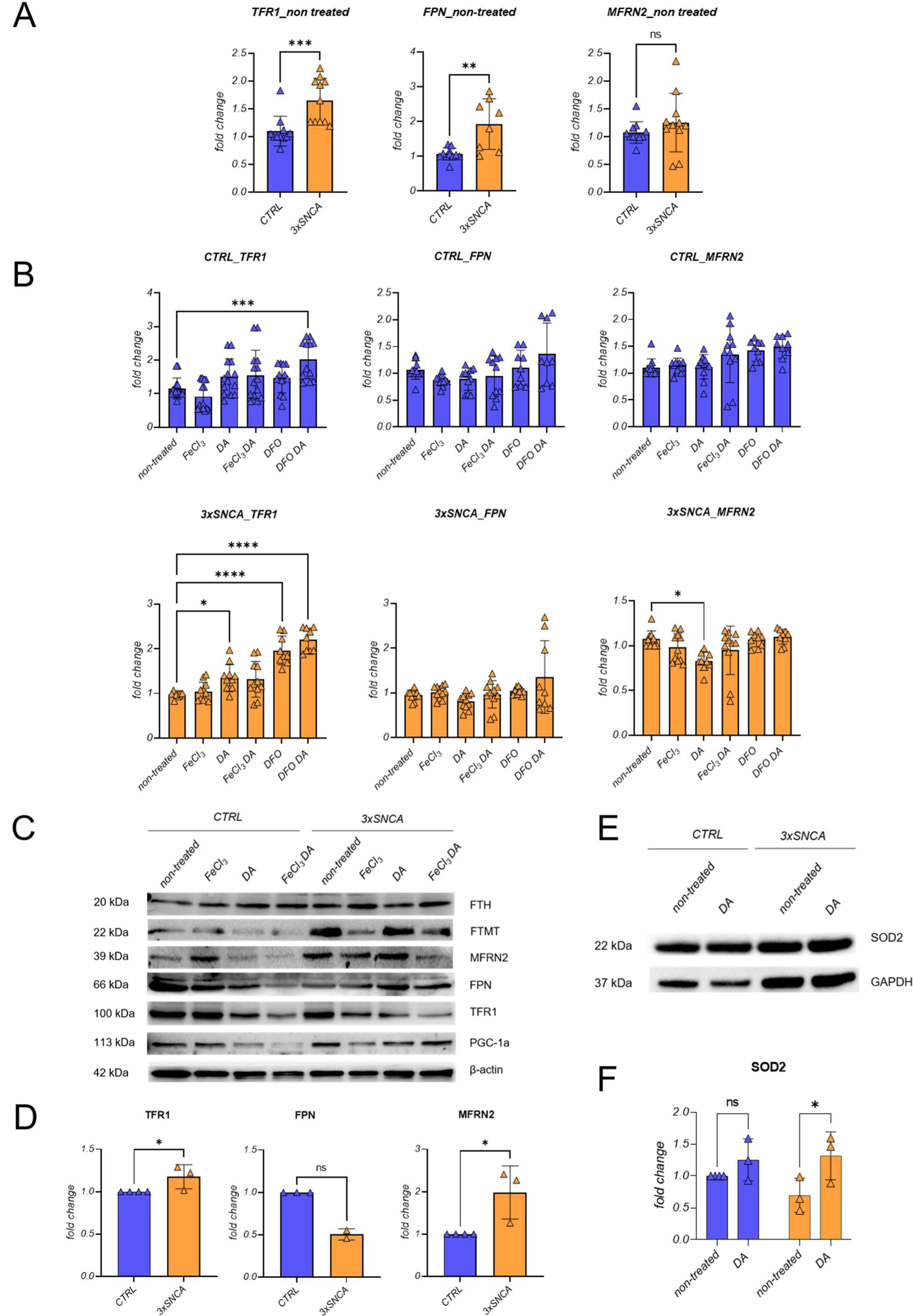
Expression of iron- and mitochondria-related genes and proteins in hiPSC-derived neurons. A) Expression of TFR1, FPN, and MFRN2 normalized with tubulin in the CTRL and PD patient (3xSNCA) lines at basal conditions. B) Impact of the compounds of interest on genes involved in cellular iron metabolism and mitochondrial iron status. C) Representative western blot showing FTH, FTMT, MFRN2, FPN TFR1, PGC-1α protein expression upon treatments. Protein levels were analyzed with the indicated antibodies, and β-actin was used as a loading control. Molecular mass markers are in kilodaltons (kDa). D) Densitometric analyses of proteins found differently expressed in hiPSC-derived neurons of a control and PD patient in untreated conditions after being resolved with WB. E) Representative western blot of SOD2 protein levels upon iron and DA treatments in CTRL and mutant cells. F) Quantification of SOD2 levels in CTRL and mutant cells. n≥2. β-actin and GAPDH were used as loading control. Statistical differences were calculated by Mann Whitney test for comparisons between the two groups in untreated conditions (A, D), ordinary one-way ANOVA followed by Dunnett’s test to correct for multiple testing was applied for comparison of multiple groups (B). For comparisons between non-treated and DA treated cells (E), a two-way ANOVA was performed followed by Šídák’s test to correct for multiple testing. *p≤0.05, ***p≤0.001; ****p≤0.0001.

Since several iron transporters are regulated post transcriptionally via iron regulatory proteins and the iron-responsive element (IRE) signaling pathway (Zhou and Tan, 2017), it is important to assess their protein levels. When comparing protein levels of non-treated control and PD patient neurons, we observed that TFR1 (p=0.029) and MFRN2 (p=0.029) protein levels were higher in the 3x*SNCA* line compared to the control line. Furthermore, contrarily to the mRNA level, FPN protein expression was lower in the PD line compared to the control (**Figure 4C, D**). This again suggests that cellular iron trafficking differs in the two lines. In particular, in the patient line, alterations are suggestive of intracellular/cytoplasmic iron deficiency as reflected by higher TFR1 and lower FPN levels compared to the control line. On the other hand, higher MFRN2 in the PD line suggests that iron is transferred to mitochondria thereby promoting mitochondrial iron accumulation. In summary, the balance between a predicted lower cytoplasmic iron content and iron accumulation in mitochondria resulted in no obvious change in total cellular iron levels as implied by the measurement of total intracellular iron content (**Figure S7**). Regarding treatments, we detected no changes of TFR1 protein levels in the control and PD lines after 16 hours of dopamine supplementation, while iron and DFO had to some extent the anticipated effects (**Figure 4**; **Figure S8**). FPN protein levels were significantly decreased by dopamine in the control line, whereas in the patient line the low FPN levels increased with dopamine treatment (**Figure 4C; Figure S8)**. MFRN2 levels as indicator of mitochondrial iron status remained largely unchanged by either treatment in both lines (**Figure S8**).

When analysing peroxisome proliferator-activated receptor gamma coactivator-1α (PGC-1α), the major regulator of mitochondrial biogenesis, in the control line, dopamine decreased its protein levels (DA: p<0.0001; FeCl_3_ DA: p<0.0001), contrarily to the PD patient line, where dopamine supplementation increased PGC-1α levels (DA: p=0.034; FeCl_3_ DA: p=0.007) (**Figure 4C; Figure S8**). We also measured superoxide dismutase 2 (SOD2) protein levels as an indicator of cellular stress responses. Dopamine increased SOD2 levels in both lines, but this was significant only in the PD patient neurons (p=0.039) (**Figure 4E, F**).

### Reduced oxygen consumption rate in hiPSC-derived patient neurons improves upon dopamine treatment

To evaluate cellular respiration in hiPSC-derived neurons (control and PD patient line) (**Figure 5A**), we quantified basal extracellular oxygen consumption rate (OCR) in ROUTINE state and after induction of maximal respiration with FCCP, in both control and patient hiPSC-derived neurons with and without dopamine and FeCl_3_ treatments (**Figure 5B**).

**Figure 5.**
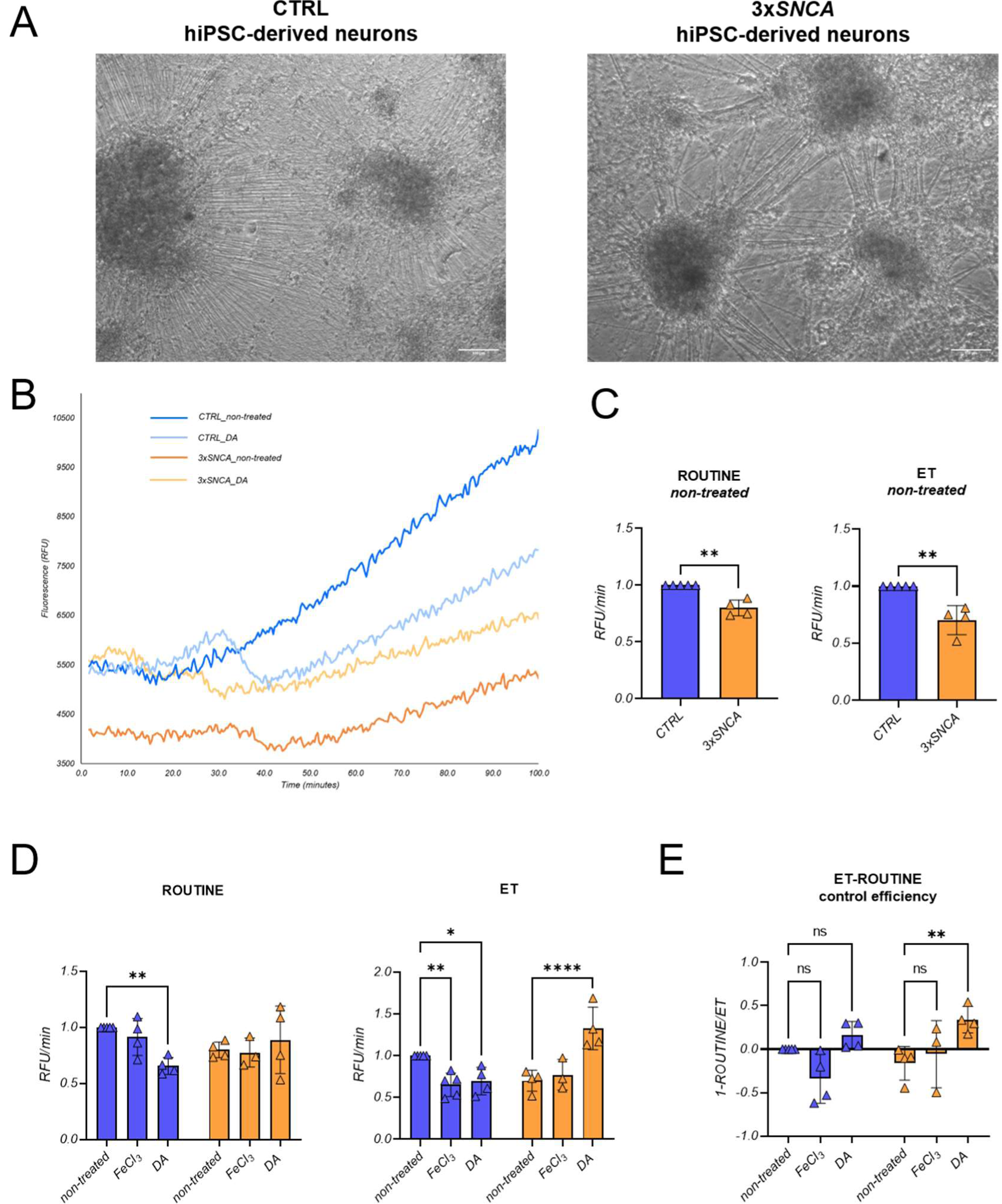
Oxygen consumption rates in hiPSC-derived neurons of control and PD patient line with 3xSNCA mutation. A) Representative light microscopy images of hiPSC-derived neurons used for OCR measurement at day 34 of differentiation. Scale bar: 100 µm. B) Representative curves of the maximal respiration fluorescence signal in hiPSC-derived neurons of the non-treated control (CTRL, non-treated, blue line) versus control cells treated with dopamine (CTRL_DA, light blue line) and non-treated 3xSNCA cells (3xSNCA, non-treated, orange line) versus 3xSNCA cells treated with dopamine (3xSNCA_DA, yellow line). C) Relative quantification of ROUTINE and maximal respiration measured after the addition of 2.5 µM FCCP in the control and the PD patient lines without treatment. D) ROUTINE and maximal respiration in the control and PD patient lines after treatments with FeCl_3_ or dopamine E) ET-ROUTINE control efficiency calculated as 1-ROUTINE/ET in control and PD patient-derived lines after treatments with FeCl_3_ or dopamine. n≥3 independent differentiations with 3 wells per respiratory parameter for each differentiation. Error bars represent the mean ±SD of at least 3 data points (one for each differentiation) for each experimental group. p values were determined using two-way ANOVA followed by Šídák’s test to correct for multiple comparisons. *p≤0.05; **p≤0.01; ***p≤0.0001; ****p≤0.00001. RFU: relative fluorescence units.

In non-treated conditions, the PD line showed a significantly decreased OCR in ROUTINE (p=0.008) and ET capacity (p=0.008) compared to the control line (**Figure 5C**). In the control line, dopamine treatment resulted in significantly reduced ROUTINE respiration compared to untreated cells (p=0.022), while ROUTINE respiration was not affected in the PD patient line (**Figure 5D)**. After induction of maximal respiration, both FeCl_3_ and dopamine treatments decreased OCR levels in the control line (FeCl_3_: p=0.007; DA: p=0.025), while in the 3x*SNCA* line, dopamine treatment increased maximal respiratory capacity to control levels (p<0.0001) (**Figure 5D**).

Additionally, we calculated ET-ROUTINE control efficiency, which is an expression of the relative scope of increasing ROUTINE respiration in living cells by uncoupler titrations to obtain ET capacity, and we found a significant increase in the PD patient line upon dopamine supplementation (p=0.007) (**Figure 5E**). Reduced mitochondrial respiration in control neurons upon dopamine treatment is in line with observations made in SH-SY5Y cells, while in patient neurons dopamine induces a beneficial effect on OCR.

### Metabolic signature in hiPSC-derived patient neurons changes upon dopamine treatment

To better understand the metabolic changes associated with observed mitochondrial alterations between control and patient neurons, we performed an untargeted metabolomic analysis in cell pellets of neurons treated with iron or dopamine for 16 hours. Both polar and non-polar metabolites were analyzed (**Figure S9; Figure S10**). Among the identified polar metabolites confirmed with standards or putatively identified by *in silico* fragmentation, 12 were significantly different between patient and control neurons under basal conditions, while 4 metabolites showed a trend towards reduction (malic acid, methionine) or increase (inosine, citrulline) in the PD line compared to the control line. Specifically, threonine, serine, glutamic acid, glutamine, taurine, N-acetylneuramic acid, uridine 5’-monophosphate, acetylcarnitine, and creatine were significantly decreased in the PD line compared to the control, while succinic acid, propionylcarnitine, and lactate were significantly increased in the PD line (**Figure S9**). In addition, we calculated the ratio between propionylcarnitine and acetylcarnitine as additional biomarker of abnormalities in propionate catabolism as suggested by increased propionyl carnitine (Collado et al., 2020), demonstrating an increased ratio in the PD line compared to the control line (**Figure S11A**). Among the investigated metabolites, those affected most by iron or dopamine treatments were threonine, serine and glutamic acid in both lines but with opposite trends: while in the control neurons dopamine resulted in significantly reduced levels, in the PD patient neurons dopamine treatment increased the levels of these metabolites towards the concentrations found in untreated control neurons (threonine – CTRL FeCl_3_: p=0.003; DA: p=0.005 – 3x*SNCA* DA: p=0.012; serine – CTRL DA: p=0.020 - 3x*SNCA* DA: p=0.003; glutamic acid – CTRL FeCl_3_: p=0.032; DA: p=0.012 – 3x*SNCA* DA: p=0.012). Furthermore, glutamine and creatine showed a non-significant trend towards increased levels after dopamine treatment in the PD patient line and reduction in the control line. Succinic acid levels were affected by dopamine only in the PD line (p=0.037) but not in the control line (**Figure 6**). Dopamine treatment also affected the ratio between propionylcarnitine and acetylcarnitine with a trend towards decreased levels in the patient line (**Figure S11B**). A list of all detected polar metabolites is reported in **Table S3**.

**Figure 6.**
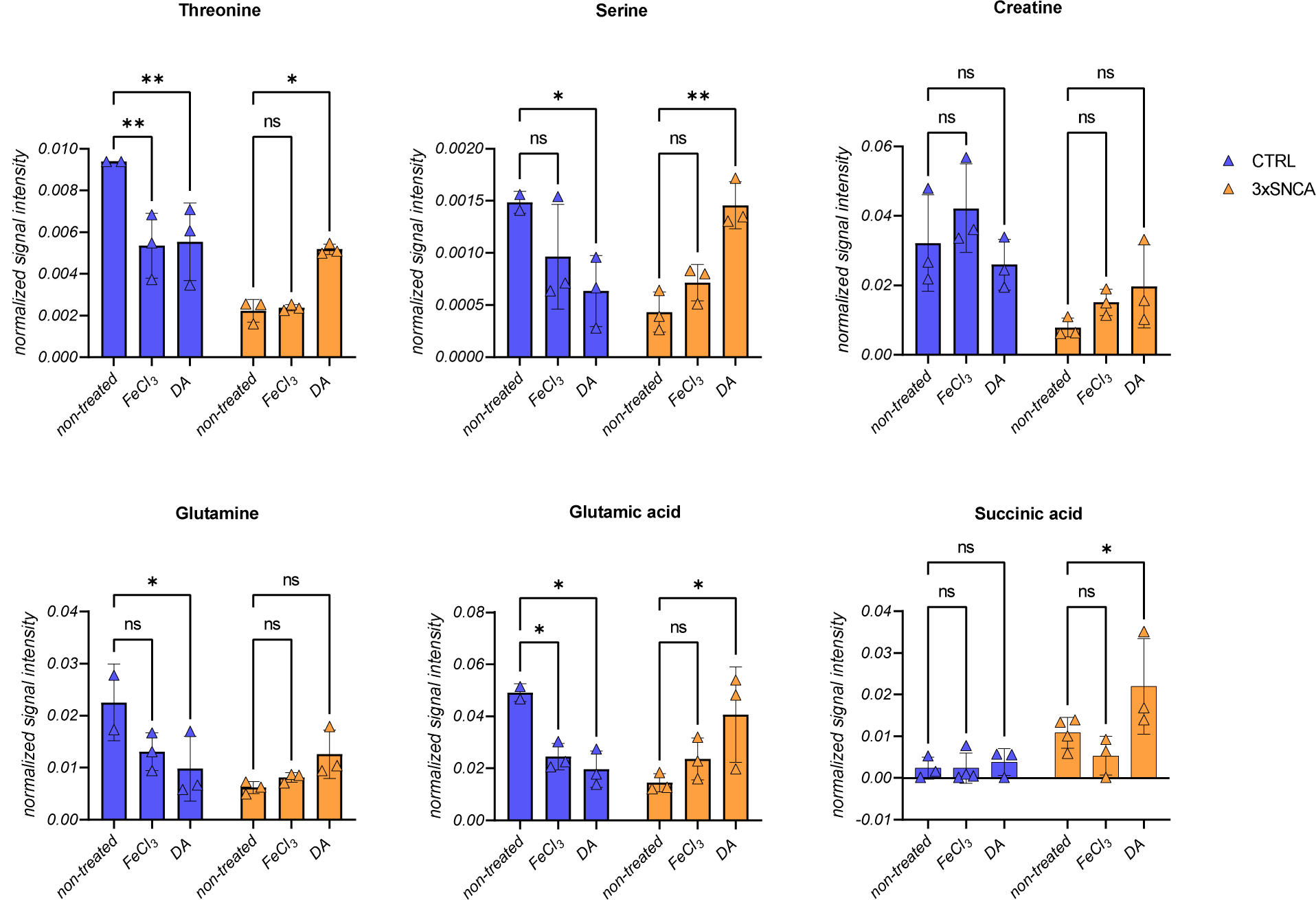
Polar metabolites affected by dopamine or FeCl_3_ treatments. Blue=Control (CTRL) line; orange=3xSNCA line. n=3 independent differentiations. Error bars represent the SD. p values were determined using a two-way ANOVA followed by Šídák’s test to correct for multiple comparisons. *p≤0.05; **p≤0.01.

Serine, threonine, and creatine belong to the glycine, serine, threonine metabolism pathway, and they are all crucially involved in mitochondrial function and energy production (**Figure 7**). Their reduction in the PD neurons compared to the control neurons (**Figure S9**) further confirms impaired energy metabolism in the PD line.

**Figure 7.**
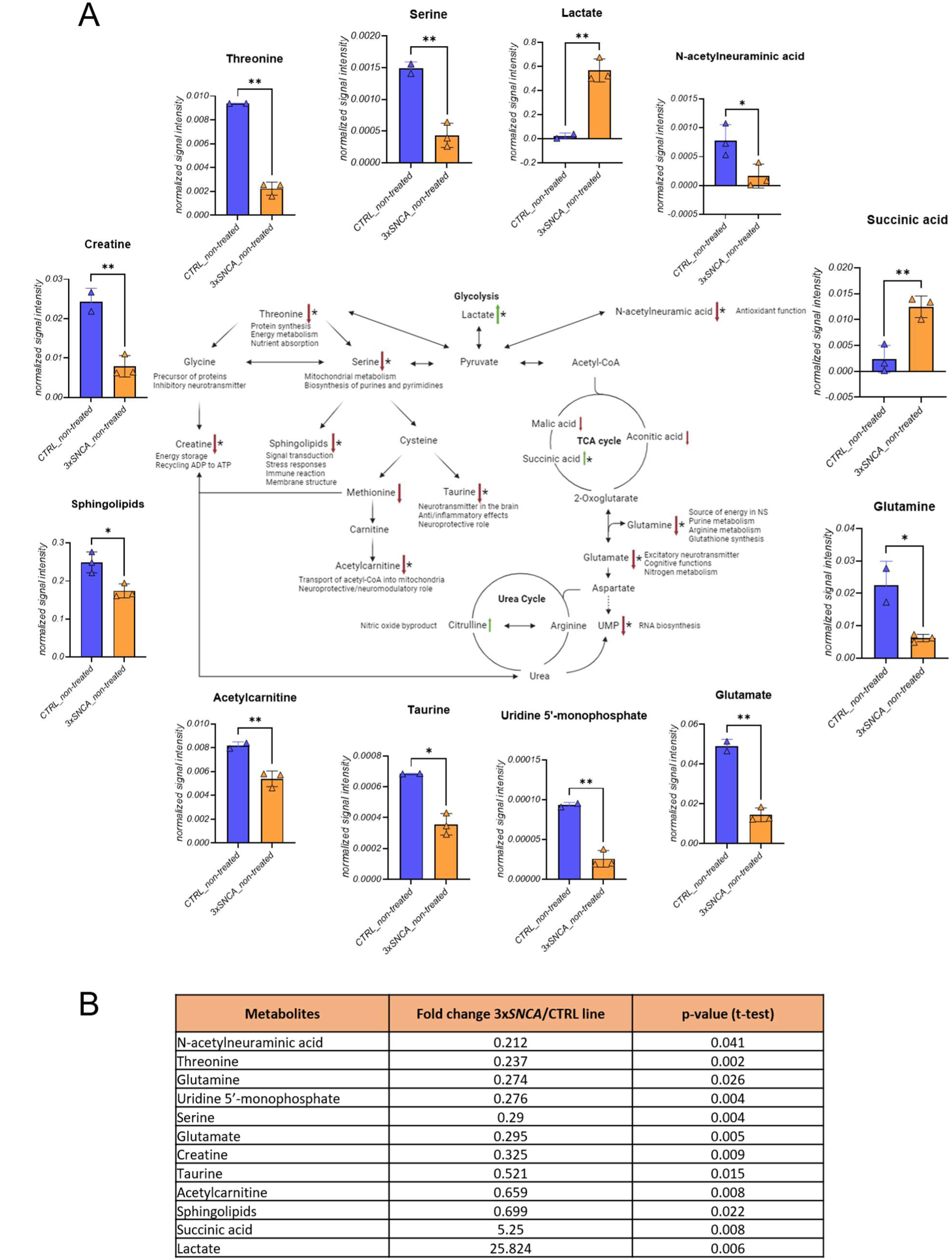
Metabolites changing between CTRL and 3xSNCA hiPSC-derived neurons and their proposed involvement in metabolic pathways. A) Detected metabolites were linked indicating bioenergetics processes and “branch points”, where changes in PD (3xSNCA) cells occur. Particularly, alterations in the pathway involving threonine, serine, creatine and acetylcarnitine may result in impaired mitochondrial functioning given their crucial role in mitochondrial activities. Similarly, reduction in Krebs cycle metabolites and downstream metabolites such as glutamine and glutamate may be associated to oxidative stress and consequent cell damage. Black arrows indicate the progression of the pathway, with double arrows indicating that the metabolite can be derived from both directions; red arrows indicate decreased metabolites, while green arrows indicate increased metabolites in the PD line compared to the control line. Bar graphs represent normalized signal intensities for selected metabolites in the control (CTRL) and PD-derived (3xSNCA) cell lines. Asterisks indicate metabolites that were significantly different between the two lines. B) Table reporting the metabolites that were significantly different between CTRL and 3xSNCA with their respective fold change (3xSNCA/CTRL) and p-values. Image created with Biorender.

Among the non-polar metabolites, 100 features were detected, of which 18 and 17 metabolites were tentatively identified at level 3 in positive and negative ionization modes, respectively (**Table S4**). Ten lipids were significantly altered in the patient hiPSC-derived neurons under basal culture conditions, including coenzyme Q10, which is a crucial electron carrier in the electron transfer system and a lipophilic antioxidant preventing the generation of free radicals, and some glycerophospholipids/sphingolipids (**Figure S11B, C**). Iron treatment did not influence these metabolites, and dopamine affected lipid levels only in the control line (i.e. PE(18:0/0:0), PC(18:1(9E)/0:0)) but not in the patient line, except for cholesterol sulfate, which was decreased upon dopamine treatment reaching levels comparable to those found in the control line (**Figure S12**). Examination of the metabolic changes by lipid class (**Figure S13A, B**) revealed that both sphingolipids and glycerophospholipids are reduced in the PD line compared to the control line, and dopamine did not affect the levels of these lipids (**Figure S13C, D**). In general, the main metabolic alterations are related to mitochondrial metabolism (**Figure 7**), and dopamine supplementation beneficially affected their dysregulated levels in PD cells.

## Discussion

In this work, we examined the linkage between iron and dopamine metabolism in SH-SY5Y neuroblastoma cells and hiPSC-derived dopaminergic neurons. We observed that dopamine affects iron homeostasis, mitochondrial respiration, and oxidative stress by altering crucial iron transporters, leading to perturbations of intracellular and mitochondrial iron levels. These dopamine-induced alterations of iron homeostasis can result in the observed alterations of mitochondrial function (Lane et al., 2015; Oexle et al., 1999). Similarly, changes of subcellular iron distribution, oxygen consumption, and mitochondrial metabolism were observed in healthy control hiPSC-derived dopaminergic neurons upon dopamine treatment. In contrast, in hiPSC-derived PD patient neurons, characterized by changes in iron homeostasis, dopamine promoted a beneficial effect by improving mitochondrial functionality.

Treatment with dopamine for 16 hours resulted in cellular damage, including decreased mitochondrial respiration and related mitochondrial alterations (reduced mtDNA copy number and citrate synthase activity), increased oxidative stress, and reduced iron-dependent aconitase activity, features observed both in SH-SY5Y neuroblastoma cells and in control hiPSC-derived neurons. Dopamine may therefore push the cells to the limits of their antioxidation/detoxification capacity when supplemented in excess. These data agree with the observation of Burbulla and colleagues, who described that Levodopa treatment, the precursor to dopamine that is used as a dopamine replacement agent in PD, was able to increase oxidized dopamine levels in human control neurons (Burbulla et al., 2017).

In SH-SY5Y cells, we found that dopamine treatment increased TFR1 expression both on mRNA and protein levels. The promotion of iron uptake induced by dopamine may be further related to a direct interaction between these two compounds, resulting in the formation of dopamine quinones and an increased cytoplasmic retention of iron as labile iron pool, more prone to oxidative events. This is in line with the chemical structure of dopamine bearing a catechol, a known iron-binding moiety (Zhang et al., 2020). Binding of iron to dopamine might result in a dysregulated iron homeostasis. Alternatively, iron might promote dopamine oxidation leading to toxic products such as aminochrome (Aguirre et al., 2012; Sun et al., 2018). In addition to these possibilities, accumulated iron can damage mitochondria via the Fenton reaction (Cerri et al., 2019; Ward et al., 2014) or by activating inflammatory molecules such as TNF-α and IL-6 (Martin-Bastida et al., 2021). Iron retention in the cytoplasm upon dopamine supplementation is further indicated by stable FTH levels and no upregulation of FPN protein levels, although it was increased on the mRNA level (Tian et al., 2020; Zhang et al., 2020).

Since dopamine affects iron levels, the function of iron-sulfur-cluster containing enzymes such as ACO1 and ACO2 may be affected due to altered iron levels or oxidative stress (Pantopoulos et al., 1997) resulting in decreased protein and activity levels of these enzymes. It is known that hydrogen peroxide or nitric oxide can disrupt the iron-sulfur cluster of ACO1 activating its iron regulatory protein 1 (IRP1) conformation. This promotes intracellular iron accumulation by increasing TFR1 and DMT1 mRNA stability (Liu et al., 2022). Furthermore, superoxide anion, which we found increased upon dopamine treatment, was shown to inactivate ACO2 (Liang and Patel, 2004), suggesting that oxidative stress may decrease the activity of this enzyme, leading to a predicted repercussion on the cellular metabolism. Alternatively, iron loading can reduce ACO2 expression via IRP1 and 2 mediated translational repression (Muckenthaler et al., 2017). Dopamine-mediated damage of mitochondria was confirmed by reduced mitochondrial quality, reduced mtDNA copy number and reduced citrate synthase activity. Indeed, excessive free radical products created during energy production and dopamine metabolism can lead to damage of both mtDNA and the overall mitochondrial machinery (Swerdlow et al., 1996; Yan et al., 2013). Even though iron administered alone showed little effects on mitochondrial function, dopamine-caused impairments on mitochondria could not be overcome by concomitant substitution of iron.

Contrarily, in pathological conditions (represented by PD patient derived hiPSC-neurons with a 3x*SNCA* mutation) dopamine treatment led to a rescue effect shown by improved respiration, activation of antioxidant pathways (shown by SOD2 levels), and a series of positive effects on metabolites involved in mitochondrial function. In a recent work it has been found that tetrahydrobiopterin (BH4) protected from genetically and chemically induced PD-related stressors in mice and human neurons from PD patients (Cronin et al., 2023). BH4 is an essential co-factor for enzymes involved in various cellular processes, for instance nitric oxide production, phenylalanine metabolism, and neurotransmitter biosynthesis, including dopamine synthesis (Werner et al., 2011). Cronin and colleagues observed that BH4 enhanced mitochondrial fitness and maintained firing rates of dopaminergic neurons (Cronin et al. 2023). In our model, comparable effects were fulfilled by dopamine administration. Furthermore, BH4 plays a role in preventing ferroptosis (Cronin et al., 2019). Likewise, dopamine may prevent ferroptosis by correcting iron imbalances in human dopaminergic neurons of PD models. Of note, a recent clinical study using the iron chelator deferiprone (DFP) to target pathological iron accumulation in patients with early onset PD failed to show clinical benefit and even resulted in worsening of symptoms (Devos et al., 2022). This points to the importance of spatio-temporal distribution of iron and dopamine for maintaining essential function.

When comparing non-treated neurons, higher TFR1 and lower FPN levels in the hiPSC-derived patient neurons suggested cytoplasmic iron deficiency opposite to evidence of mitochondrial iron accumulation, as reflected by higher MFRN2. However, while the combination of dopamine with FeCl_3_ significantly decreased TFR1 levels in the control line, treatments in the patient line – at least after 16 hours – did not affect TFR1 protein levels. This is in line with a shift of iron from the cytoplasm to mitochondria thereby prohibiting a prolonged increase of cytoplasmic iron levels, which could affect TFR expression via IRP/IRE interaction. Significant variations of FPN and FTH only in the control line after treatments further suggested that the two lines responded differently to perturbations of iron homeostasis. Failure in storage regulation with consequently increased levels of labile iron, as opposed to ferritin-bound iron, is known as a possible cause of iron-related cellular dysfunctions in PD (Tian et al., 2020). If free iron content is elevated, the Fenton reaction can generate ROS and therefore promote oxidative damage (Cheng et al., 2022). Accordingly, we found that SOD2 levels responded to dopamine treatment by alleviating mitochondrial oxidative stress in our PD model. Our data further suggested impaired iron handling in the PD neurons, in which mitochondrial iron overload may be prevented by dopamine treatment, for example through shuttling excess iron out of mitochondria. Moreover, in line with well-established mitochondrial defects, we measured decreased respiration in the 3x*SNCA* patient neurons compared to the control neurons. Multiple biological mechanisms may underlie this phenotype including a possible accumulation of α-syn in the PD neurons, increased amounts of oxidized dopamine (Burbulla et al., 2017), or iron overload (Fischer et al., 2021; Huang et al., 2018). Physiological responses to impaired mitochondria include a compensatory increase of transcription of PGC-1α, the major regulator of mitochondrial biogenesis (Lin et al., 2009). In contrast to control neurons, treatment of patient neurons with dopamine increased PGC-1α, underscoring again the beneficial effect of dopamine treatment in the PD model, which is in line with the observation of ameliorated mitochondrial function in restless legs syndrome subjects receiving Levodopa (Haschka et al., 2019).

Of importance, an untargeted metabolomics approach revealed that most of the observed changes in the metabolism between the control and the PD patient model are attributable to energy/mitochondrial metabolism (e.g., threonine, serine pathways) and antioxidant mechanisms (e.g., glutamine). Interestingly, dopamine increased these metabolites in the PD line, while disturbing their balance in the control, pointing to a damaging effect of dopamine in control neurons and a beneficial effect only in patient neurons. Glutamine is the precursor of glutamate and glutathione, the principal excitatory neurotransmitter and a crucial antioxidant in the body (Lau and Tymianski, 2010; Zhang et al., 2019). The effect of dopamine on increasing these metabolite levels in the PD neurons but not in control neurons might be an indication of a compensatory effect to underlying intrinsic oxidative stress. Instead, reduced levels of glutamine and glutamate in baseline conditions in the PD line may indicate their consumption to produce glutathione, and therefore to neutralize oxidative stress. Several TCA metabolites, including malic acid and aconitic acid, were found reduced in PD neurons compared to the control cells, being in line with the previously observed reduction in TCA cycle metabolites in PD (Shao and Le, 2019). Succinic acid was found increased as in (Zambon et al., 2019). Beyond its production in TCA, succinate is also produced from the degradation of odd-chain fatty acids, methionine, threonine, leucine, or isoleucine. All of these metabolites are finally converted to succinate with propionyl-CoA, methylmalonyl-CoA and succinyl-CoA as intermediates (Collado et al., 2020). Along with decreased levels of these precursors (methionine, threonine), we also observed an increase of propionylcarnitine and the ratio propionylcarnitine/acetylcarnitine in the patient line compared to the control line. Propionylcarnitine accumulation might indicate abnormalities in propionate catabolism or its excess production (confirmed also by its ratio), where excess propionate is transferred to carnitine to reduce its toxicity (Collado et al., 2020; Wongkittichote et al., 2019). In previous studies, short-chain fatty acids (SCFA) and the propionate pathway have been found to be dysregulated in PD (Chen et al., 2022; He et al., 2021; Levin et al., 2010; Park et al., 2017; Qi et al., 2023). Although there is controversy about the content of acetic acid and propionic acid (Qi et al., 2023), and these studies are conducted on serum or stool, increased levels of SCFAs, including propionic acid, were found in serum of PD patients compared to controls (Chen et al., 2022), confirmed also previously observed results (He et al., 2021). Interestingly, plasma levels of propionic acid were associated to more severe motor symptoms (Chen et al., 2022). Moreover, elevated levels of methylmalonate and homocysteine were observed in PD (Levin et al., 2010) suggesting that both might be markers for neurodegenerative processes, and both are linked to propionate metabolism. Furthermore, in serum, methylmalonic acid correlated with neuropathic pain in PD (Park et al., 2017) and low vitamin B12 levels, which is a master regulator of this pathway, are associated with a worse prognosis of PD, demyelination and peripheral neuropathy (Cardoso, 2018). In summary, dopamine treatment induced an opposite effect in the control neurons compared to the PD patient neurons only for some of the analyzed metabolites (e.g., threonine, serine, and glutamine pathway), suggesting that dopamine may have an impact on those specific metabolic pathways.

Even though this work shows an important role for the dopamine-iron interplay in neuronal cell models, additional aspects of the biological mechanisms underlying the rescue of PD patient cells in relation to iron homeostasis may still need to be clarified. For instance, we measured protein expression at a single time point and may thus have missed early changes or detected the sum of dopamine-mediated alterations and counter regulatory signals. In addition, the study of co-culture/organoids of dopaminergic neurons with other neuronal populations or *in vivo* studies might help to reconstitute a more physiological environment.

### Conclusion

In summary, under physiological conditions and pH levels, dopamine may constitute a complex with iron, which can be delivered across membranes (Goetz et al., 2002) and result in the formation of oxygen radical species through chemical reactions (Arreguin et al., 2009). In PD, iron regulation is already impaired due to intrinsic PD pathogenic mechanisms such as presence of increased oxidative stress, as well as misfolded α-syn (Arreguin et al., 2009). Therefore, administration of dopamine may partially rescue a system characterized by increased mitochondrial iron levels and importantly, it appears to act via mitochondrial processes. Indeed, dopamine can increase impaired oxygen consumption rate, upregulate PGC1α and reduce ROS production by regulating the expression and activity of ROS detoxifying enzymes. Dopamine may be able to shuttle iron across membranes and between cytoplasm and mitochondria, thus reconstituting altered intracellular iron distribution. At early stages of the disease, the correct balance of both iron and dopamine may play a crucial role in vulnerable cells, such as dopaminergic neurons, and the minimum misbalance of these delicate compounds together with external (e.g., environmental) or internal (e.g., genetic) factors may corrupt the metabolic and proteostatic equilibrium and result in disease. Contrarily, at later stages of the disease, when neurons are already damaged and show a deficiency in endogenous dopamine production, treatment with dopamine can restore some of the metabolic alterations of dopaminergic neurons and improve mitochondrial function. A further dissection of this pathway and its involvement in the pathogenesis of PD might be useful for the development of therapeutic approaches that can restore altered intracellular iron distribution and prevent the formation of nitrating/oxidative agents as a means of limiting neuronal injury in PD.

## DECLARATIONS

### Ethics approval and consent to participate

The study was approved by the Ethics Committee of the Healthcare System of the Autonomous Province of Bozen/Bolzano (approval number 102-2014 dated 26.11.2014 with extension from 19.02.2020). Study participants provided their written informed consent to participate in this study.

### Consent for publication

Study participants have given their consent for publication in an anonymous form.

### Availability of data and material

All data generated during this study are included in this article and its supplementary information file. As for the metabolomics data, the informed consent provided by the study participants does not allow the upload of individual-level metabolomic data to public repositories.

## Authors’ contributions

CB: designed and performed experiments, analyzed the data and wrote the manuscript; MS: designed and performed experiments, reviewed the manuscript; ML: designed and performed experiments, analyzed the data, participated in the interpretation of results, wrote and revised the manuscript; CMG: provided support for the metabolomic analyses, revised the manuscript; BTA: provided support for the metabolomic analyses, revised the manuscript; MPCR: designed and performed experiments, revised the manuscript; CF: helped with experiments and helped revise the manuscript; CD: provided support for Oxygraph experiments and revised the manuscript; HT: performed the atomic absorption spectroscopy experiments; AZ: designed and performed experiments, revised the manuscript; PPP: provided critical review of the manuscript; ELS: performed critical input and revised the manuscript; IP: designed and supervised the project, participated in the interpretation of results, wrote and revised the manuscript; GW: designed and supervised the project, provided the interpretation of results, wrote and revised the manuscript.

All authors read and approved the final manuscript.

## Acknowledgements

The authors acknowledge the metabolomics platform of the LCSB and Lorenzo Favilli for the support during the LC-MS experiments and analysis. The authors also thank Giovanna Gentile, Valentina Gilmozzi, and Chiara Volani for experimental support and Gianfranco Frigerio for his help hosting the exchange at LCSB. CD has been employed by Oroboros Instruments.

