## Supplementary Material for "Dopamine-iron homeostasis interaction rescues mitochondrial fitness in Parkinson’s disease"

Chiara Buoso[a,b], Markus Seifert[b,c], Martin Lang[a], Corey M. Griffith[d], Begoña Talavera Andújar[d], Maria Paulina Castelo Rueda[a], Christine Fischer[b], Carolina Doerrier[f], Heribert Talasz[e], Alessandra Zanon[a], Peter P. Pramstaller[a], Emma L. Schymanski[d], Irene Pichler[a*], Guenter Weiss[b,c*]

[a] *Institute for Biomedicine, Eurac Research, Affiliated Institute of the University of Lübeck, Bolzano, Italy.*

[b] *Department of Internal Medicine II, Medical University of Innsbruck, Innsbruck, Austria.*

[c] *Christian Doppler Laboratory for Iron Metabolism and Anemia Research, Medical University of Innsbruck, Innsbruck, Austria.*

[d] *Luxembourg Centre for Systems Biomedicine (LCSB), University of Luxembourg, Belvaux, Luxembourg.*

[e] *Institute of Medical Biochemistry, Protein Core Facility, Biocenter Innsbruck, Medical University of Innsbruck, 6020 Innsbruck, Austria.*

[f] *Oroboros Instruments, 6020 Innsbruck, Austria.*

*These authors have contributed equally

***S1 Methods***

*S1.1 Extraction of intracellular metabolites*

During the metabolite extraction, the following internal standards (IS) were used: ([UL-13C5]Ribitol, 2 µg/ml; pentanedioic-d6 acid, 2 µg/ml; tridecanoic-d25 acid in 0.9% NaCl (w/v) solution, 10 µg/ml; 6-chloropurine riboside in MilliQ water, 10 µg/ml; 4-chloro-DL-phenylalanine in aq. 0.1 M HCl, 10 µg/ml; Nε-trifluoroacetyl-L-lysine, in aq. 0.1 M HCl, 10 µg/ml; thionicotinamide adenine dinucleotide in MilliQ water, 10 µg/ml). Information on the standards included in the polar mix used during the HILIC runs is given in **Table S1.**

*S1.2 Modifications for LC-HRMS analysis (non-polar extracts)*

Non-target liquid chromatography high resolution mass spectrometry (LC-HRMS) analysis was performed with a Q Exactive Orbitrap using both hydrophilic interaction liquid chromatography (HILIC) and reverse phase (RP) columns. The samples were analyzed in both positive (+) and negative (−) electrospray ionization (ESI) modes. A polar mix was included in the worklist and injected during the run (**Table S1**). Quality control (QC) included the use of internal standards (IS); blank injections were made every 5-10 samples. IS retention times (RT) were checked throughout the runs to assess the consistency of the results.

For non-polar extracts, an Acquity UPLC CSH C18 RP column (100 × 2.1 mm; 1.7 µm) was used. The method was adapted from (Cajka, Smilowitz, and Fiehn 2017). Briefly, mobile phase (MP) A was 60:40 (v/v) ACN:MilliQ H2O with ammonium formate (10 mM) and formic acid (FA) (0.1%) while MP B was 90:10 (v/v) isopropanol:ACN with ammonium formate (10 mM) and FA (0.1%). The gradient program was 85A/15B at 0 min, 85/15 at 2 min, 52/48 at 2.5 min, 18/82 at 11 min, 1/99 at 15 min, 85/15 at 15.1 min, and 85/15 at 19 min. The column temperature was set at 65°C and the flow rate at 0.6 mL/min. The sample injection volume was 5 µl. The following full MS/data dependent (dd) MS2 settings were employed: scan range *m/z*=60–900 in ESI (+), and *m/z* =120-1200 in ESI (−); resolving power 120,000 FWHM (*m/z* 200) in ESI (+) and 60,000 FWHM (*m/z* 200) in ESI (−); maximum injection time (IT) 70ms and 100ms for ESI (+) and ESI (-), respectively; and automatic gain control (AGC) target was 1.0 $\times 10^6$. For the dd-MS2/dd-SIM (data-dependent selected ion monitoring), the following settings were used: resolving power 30,000 FWHM (*m/z* 200) for ESI (+) and 15,000 (*m/z* 200) for ESI (−); loop count 5 for ESI (+) and 4 for ESI (-); Top N 5 for ESI (+) and 4 for ESI (-); maximum IT 70ms for ESI (+) and 50ms for ESI (-); and normalized collision energy, 30% in ESI (+) and 20%, 30% and 40% for ESI (−).

For polar extracts, a SeQuant® ZIC-pHILIC 5-µm polymer (150×2.1 mm) column was used. LC and MS parameters were previously described (Talavera Andujar et al. 2022). Briefly, MP A was 20 mM ammonium acetate in ACN, while MP B was 20 mM ammonium acetate in MilliQ $H_2O$. The HRMS was operated in full scan profile mode with a scan range of 60-900 *m/z*.

### *S2 Results*

*S2.1 Supplementary tables*

***Table S1***. *List of metabolites included in the standard mix. Monoisotopic mass refers to the neutral mass. $[M\text{-}H]^-$ refers to a negatively charged molecular ion (resulting from the loss of an hydrogen ion) while $[M+H]^+$ refers to a positively charged molecular ion (formed after the addition of a proton).*

| HMDB Name | Reference | HMDB ID | Monoisotopic mass | $[M\text{-}H]^-$ | $[M+H]^+$ |
|---|---|---|---|---|---|
| 2-Aminobenzoic acid | A89855-25G | HMDB0001123 | 137.0477 | 136.0399 | 138.0555 |
| 2-Hydroxyadipic acid | 92826-10MG | HMDB0000321 | 162.0528 | 161.0450 | 163.0606 |
| 2-Hydroxybutyric acid | 220116-5G | HMDB0000008 | 104.0473 | 103.0395 | 105.0551 |
| 2-Ketobutyric acid | K401-10G | HMDB0000005 | 102.0317 | 101.0239 | 103.0395 |
| 2-Methylglutaric acid | 129860-5G | HMDB0000422 | 146.0579 | 145.0501 | 147.0657 |
| 3,4-Dihydroxybenzeneacetic acid | 850217-1G | HMDB0001336 | 168.0423 | 167.0345 | 169.0501 |
| 3-Hydroxyanthranilic acid | 148776-250MG | HMDB0001476 | 153.0426 | 152.0348 | 154.0504 |
| 3-Hydroxybutyric acid | 166898-1G | HMDB0000357 | 104.0473 | 103.0395 | 105.0551 |
| 3-Indolebutyric acid | 45532-250MG | HMDB0002096 | 203.0946 | 202.0868 | 204.1024 |
| 3-Indoleglyoxylic acid | 220019-1G | HMDB0242143 | 189.0426 | 188.0348 | 190.0504 |
| Indole-3-propionic acid | 220027-1G | HMDB0002302 | 189.0790 | 188.0712 | 190.0868 |
| 3-Methylindole | 90961-100MG | HMDB0000466 | 131.0735 | 130.0657 | 132.0813 |
| 4-Hydroxy-2-oxoglutaric acid | 96599-50MG | HMDB0002070 | 162.0164 | 161.0086 | 163.0242 |
| 4-Hydroxy-L-glutamic acid | 76157-10MG | HMDB0002273 | 163.0481 | 162.0403 | 164.0559 |
| 4-Hydroxyproline | 56250-5G | HMDB0000725 | 131.0582 | 130.0504 | 132.0660 |
| 5-Hydroxyindoleacetic acid | H8876-100MG | HMDB0000763 | 191.0582 | 190.0504 | 192.0660 |
| 5-Hydroxy-L-tryptophan | H9772-100MG | HMDB0000472 | 220.0848 | 219.0770 | 221.0926 |
| 5-Methyl-DL-Tryptophan | M0534-250MG | HMDB0254563 | 218.1055 | 217.0977 | 219.1133 |
| 6-Phosphogluconic acid | P7877-100MG | HMDB0001316 | 276.0246 | 275.0168 | 277.0324 |
| Acetyl-CoA | A2056-25MG | HMDB0001206 | 809.1258 | 808.1180 | 810.1336 |
| Acetylcysteine | A7250-5G | HMDB0001890 | 163.0303 | 162.0225 | 164.0381 |
| Adenine | A2786-5G | HMDB0000034 | 135.0545 | 134.0467 | 136.0623 |
| Adenosine monophosphate | A2252-5G | HMDB0000045 | 347.0631 | 346.0553 | 348.0709 |
| Adenosine triphosphate | A2383-5G | HMDB0000538 | 506.9957 | 505.9879 | 508.0035 |
| ADP | A2754-1G | HMDB0001341 | 427.0294 | 426.0216 | 428.0372 |
| AICAR | A9978 | HMDB0001517 | 338.0628 | 337.0550 | 339.0706 |
| 3-Hydroxyglutaric acid | TRC-H942580-10 | HMDB0000428 | 148.0372 | 147.0294 | 149.0450 |
| Betaine | B2629-50G | HMDB0000043 | 118.0868 | 117.0790 | 119.0946 |
| Biotin | B4501-100MG | HMDB0000030 | 244.0882 | 243.0804 | 245.0960 |
| CDP | C9755-25MG | HMDB0001546 | 403.0182 | 402.0104 | 404.0260 |
| Choline | C7527-100G | HMDB0000097 | 104.1075 | 103.0997 | 105.1153 |
| cis-Aconitic acid | A3412-1G | HMDB0000072 | 174.0164 | 173.0086 | 175.0242 |
| Citramalic acid | 27455-1G-F | HMDB0000426 | 148.0372 | 147.0294 | 149.0450 |
| Citric acid | C1909-500G | HMDB0000094 | 192.0270 | 191.0192 | 193.0348 |
| Creatine | C3630-100G | HMDB0000064 | 131.0695 | 130.0617 | 132.0773 |
| Creatinine | C4255-25G | HMDB0000562 | 113.0589 | 112.0511 | 114.0667 |
| Mesaconic acid | 131040-10G | HMDB0000749 | 130.0266 | 129.0188 | 131.0344 |
| Cytidine monophosphate | C1006-500MG | HMDB0000095 | 323.0519 | 322.0441 | 324.0597 |
| Cytidine triphosphate | C1506-10MG | HMDB0000082 | 482.9845 | 481.9767 | 483.9923 |
| Cytosine | C3506-1G | HMDB0000630 | 111.0433 | 110.0355 | 112.0511 |

| HMDB Name | Reference | HMDB ID | Monoisotopic mass | [M-H]- | [M+H]+ |
|---|---|---|---|---|---|
| D-2-Hydroxyglutaric acid | H8378-250MG | HMDB0000606 | 148.0372 | 147.0294 | 149.0450 |
| Deoxyadenosine triphosphate | 11934511001-100mM | HMDB0001532 | 491.0008 | 489.9930 | 492.0086 |
| D-Erythrose 4-phosphate | E0377-25MG | HMDB0001321 | 200.0086 | 199.0008 | 201.0164 |
| Desaminotyrosine | H52406-5G | HMDB0002199 | 166.0630 | 165.0552 | 167.0708 |
| D-Fructose | F0127-100G | HMDB0000660 | 180.0634 | 179.0556 | 181.0712 |
| D-Galactose | G5388-500G | HMDB0000143 | 180.0634 | 179.0556 | 181.0712 |
| D-Glucose | G8270-100G | HMDB0000122 | 180.0634 | 179.0556 | 181.0712 |
| D-Glucuronic acid | 8.14723.0010-10G | HMDB0000127 | 194.0427 | 193.0349 | 195.0505 |
| D-Glycerate 3-phosphate | P8877-10MG | HMDB0060180 | 185.9929 | 184.9851 | 187.0007 |
| Phosphoserine | P0878-1G | HMDB0000272 | 185.0089 | 184.0011 | 186.0167 |
| D-Mannose | M1134-500MG | HMDB0000169 | 180.0634 | 179.0556 | 181.0712 |
| D-Mannose 1-phosphate | M1755-25MG | HMDB0006330 | 260.0297 | 259.0219 | 261.0375 |
| D-Ribose 5-phosphate | R7750-250MG | HMDB0001548 | 230.0192 | 229.0114 | 231.0270 |
| D-Sedoheptulose 7-phosphate | MS07457 | HMDB0001068 | 290.0403 | 289.0325 | 291.0481 |
| dUMP | D3876-100MG | HMDB0001409 | 308.0410 | 307.0332 | 309.0488 |
| FAD | F6625 | HMDB0001248 | 785.1571 | 784.1493 | 786.1649 |
| Flavin Mononucleotide | F2253-25MG | HMDB0001520 | 456.1046 | 455.0968 | 457.1124 |
| Fructose 1,6-bisphosphate | F6803-10MG | HMDB0001058 | 339.9960 | 338.9882 | 341.0038 |
| O-Acetylserine | CDS020792 | HMDB0003011 | 147.0532 | 146.0454 | 148.0610 |
| Fructose 6-phosphate | F3627-100MG | HMDB0000124 | 260.0297 | 259.0219 | 261.0375 |
| Fumaric acid | 47910-1KG | HMDB0000134 | 116.0110 | 115.0032 | 117.0188 |
| Galactaric acid | M89617-100G | HMDB0000639 | 210.0376 | 209.0298 | 211.0454 |
| Galactose 1-phosphate | 345646-100MG | HMDB0000645 | 260.0297 | 259.0219 | 261.0375 |
| Galacturonic acid | 48280-5G-F | HMDB0002545 | 194.0427 | 193.0349 | 195.0505 |
| Gamma Glutamylglutamic acid | G3640-25MG | HMDB0011737 | 276.0958 | 275.0880 | 277.1036 |
| Gamma-Glutamylcysteine | G0903-25MG | HMDB0001049 | 250.0623 | 249.0545 | 251.0701 |
| Glucaric acid | 21236-25G | HMDB0000663 | 210.0376 | 209.0298 | 211.0454 |
| Gluconic acid | G9005-10mg | HMDB0000625 | 196.0583 | 195.0505 | 197.0661 |
| Glucose 1-phosphate | G9380-100MG | HMDB0001586 | 260.0297 | 259.0219 | 261.0375 |
| Glucose 6-phosphate | G7375-1G | HMDB0001401 | 260.0297 | 259.0219 | 261.0375 |
| Glutaric acid | 07438-100ML | HMDB0000661 | 132.0423 | 131.0345 | 133.0501 |
| Glutathione | G6013-5G | HMDB0000125 | 307.0838 | 306.0760 | 308.0916 |
| Glyceric acid | 61786-10MG | HMDB0000139 | 106.0266 | 105.0188 | 107.0344 |
| Glycerol 3-phosphate | 94124-10MG | HMDB0000126 | 172.0137 | 171.0059 | 173.0215 |
| Glycine | 50046-50G | HMDB0000123 | 75.0320 | 74.0242 | 76.0398 |
| Guanine | 51020 | HMDB0000132 | 151.0494 | 150.0416 | 152.0572 |
| Guanosine diphosphate | G7127-100MG | HMDB0001201 | 443.0243 | 442.0165 | 444.0321 |
| Guanosine diphosphate mannose | G5131-10MG | HMDB0001163 | 605.0772 | 604.0694 | 606.0850 |
| Guanosine monophosphate | G8377-500MG | HMDB0001397 | 363.0580 | 362.0502 | 364.0658 |
| Guanosine triphosphate | G8877-25MG | HMDB0001273 | 522.9907 | 521.9829 | 523.9985 |
| Homocysteine | H4628-1G | HMDB0000742 | 135.0354 | 134.0276 | 136.0432 |
| Hydroxykynurenine | H1771-25MG | HMDB0000732 | 224.0797 | 223.0719 | 225.0875 |
| Hydroxypropanedioic acid | 86320-10G | HMDB0035227 | 120.0059 | 118.9981 | 121.0137 |
| Indole | I3408-25G | HMDB0000738 | 117.0578 | 116.0500 | 118.0656 |
| Indole-3-acetamide | 286281-1G | HMDB0029739 | 174.0793 | 173.0715 | 175.0871 |
| Indole-3-carboxaldehyde | 129445-5G | HMDB0029737 | 145.0528 | 144.0450 | 146.0606 |

| HMDB Name | Reference | HMDB ID | Monoisotopic mass | $[M-H]^-$ | $[M+H]^+$ |
|---|---|---|---|---|---|
| Indole-3-carboxylic acid | 284734-1G | HMDB0003320 | 161.0477 | 160.0399 | 162.0555 |
| Indoleacetaldehyde | I1000-25MG | HMDB0001190 | 159.0684 | 158.0606 | 160.0762 |
| Indoleacetic acid | I2886-5G | HMDB0000197 | 175.0633 | 174.0555 | 176.0711 |
| Indoleacrylic acid | I2273-1G | HMDB0000734 | 187.0633 | 186.0555 | 188.0711 |
| Indolelactic acid | I5508-250MG-A | HMDB0000671 | 205.0739 | 204.0661 | 206.0817 |
| Indolepyruvate | I7017-1G | HMDB0060484 | 203.0582 | 202.0504 | 204.0660 |
| Inosine | I4125-5G | HMDB0000195 | 268.0808 | 267.0730 | 269.0886 |
| Inosinic acid | 57510-5G | HMDB0000175 | 348.0471 | 347.0393 | 349.0549 |
| Isocitric acid | I1252-1G | HMDB0000193 | 192.0270 | 191.0192 | 193.0348 |
| Itaconic acid | I29204-100G | HMDB0002092 | 130.0266 | 129.0188 | 131.0344 |
| Ketoleucine | 68255-1G | HMDB0000695 | 130.0630 | 129.0552 | 131.0708 |
| Kynurenic acid | K3375-250MG | HMDB0000715 | 189.0426 | 188.0348 | 190.0504 |
| L-Acetylcarnitine | A6706-1G | HMDB0000201 | 203.1158 | 202.1080 | 204.1236 |
| L-Alanine | 05130-25G | HMDB0000161 | 89.0477 | 88.0399 | 90.0555 |
| L-Arginine | A8094-100G | HMDB0000517 | 174.1117 | 173.1039 | 175.1195 |
| L-Asparagine | A0884-25G | HMDB0000168 | 132.0535 | 131.0457 | 133.0613 |
| L-Aspartic acid | A7219-100G | HMDB0000191 | 133.0375 | 132.0297 | 134.0453 |
| L-Cystathionine | C3633-100MG | HMDB0000099 | 222.0674 | 221.0596 | 223.0752 |
| L-Cysteine | 30120-10G | HMDB0000574 | 121.0197 | 120.0119 | 122.0275 |
| L-Cystine | C6727-25G | HMDB0000192 | 240.0238 | 239.0160 | 241.0316 |
| L-Glutamic acid | 49449-100G | HMDB0000148 | 147.0532 | 146.0454 | 148.0610 |
| L-Glutamine | G8540-25G | HMDB0000641 | 146.0691 | 145.0613 | 147.0769 |
| L-Histidine | H5659-25G | HMDB0000177 | 155.0695 | 154.0617 | 156.0773 |
| L-Isoleucine | I7403-25G | HMDB0000172 | 131.0946 | 130.0868 | 132.1024 |
| L-Kynurenine | K8625-25MG | HMDB0000684 | 208.0848 | 207.0770 | 209.0926 |
| L-Lactic acid | L7022-5G | HMDB0000190 | 90.0317 | 89.0239 | 91.0395 |
| L-Leucine | L8912-25G | HMDB0000687 | 131.0946 | 130.0868 | 132.1024 |
| L-Lysine | 62929-100G-F | HMDB0000182 | 146.1055 | 145.0977 | 147.1133 |
| L-Malic acid | M1000-100G | HMDB0000156 | 134.0215 | 133.0137 | 135.0293 |
| L-Methionine | 64319-25G-F | HMDB0000696 | 149.0510 | 148.0432 | 150.0588 |
| L-Phenylalanine | P5482-25G | HMDB0000159 | 165.0790 | 164.0712 | 166.0868 |
| L-Proline | P5607-25G | HMDB0000162 | 115.0633 | 114.0555 | 116.0711 |
| L-Serine | S4500-1G | HMDB0000187 | 105.0426 | 104.0348 | 106.0504 |
| L-Threonine | T8441-25G | HMDB0000167 | 119.0582 | 118.0504 | 120.0660 |
| L-Tryptophan | T8941-25G | HMDB0000929 | 204.0899 | 203.0821 | 205.0977 |
| L-Tyrosine | T8566-25G | HMDB0000158 | 181.0739 | 180.0661 | 182.0817 |
| L-Valine | V0500-1G | HMDB0000883 | 117.0790 | 116.0712 | 118.0868 |
| Maleic acid | A14596 | HMDB0000176 | 116.0110 | 115.0032 | 117.0188 |
| Malonic acid | M1296 | HMDB0000691 | 104.0110 | 103.0032 | 105.0188 |
| Melatonin | PHR1767-500MG | HMDB0001389 | 232.1212 | 231.1134 | 233.1290 |
| Methylmalonic acid | 4979 | HMDB0000202 | 118.0266 | 117.0188 | 119.0344 |
| Myo-inositol 1-phosphate | 10007777 | HMDB0000213 | 260.0297 | 259.0219 | 261.0375 |
| N-Acetylglutamic acid | 855642-25G | HMDB0001138 | 189.0637 | 188.0559 | 190.0715 |
| N-Acetyl-L-aspartic acid | 00920-5G | HMDB0000812 | 175.0481 | 174.0403 | 176.0559 |
| N-Acetylserotonin | A1824-100MG | HMDB0001238 | 218.1055 | 217.0977 | 219.1133 |
| NAD | N0632-1G | HMDB0000902 | 663.1091 | 662.1013 | 664.1169 |

| HMDB Name | Reference | HMDB ID | Monoisotopic mass | $[M-H]^-$ | $[M+H]^+$ |
|---|---|---|---|---|---|
| NADH | 10128023001 | HMDB0001487 | 665.1248 | 664.1170 | 666.1326 |
| NADP | 10128031001 | HMDB0000217 | 743.0755 | 742.0677 | 744.0833 |
| NADPH | 10107824001 | HMDB0000221 | 745.0911 | 744.0833 | 746.0989 |
| N-Formyl-L-methionine | S601047-250MG | HMDB0001015 | 177.0460 | 176.0382 | 178.0538 |
| Niacinamide | N0636-100G | HMDB0001406 | 122.0480 | 121.0402 | 123.0558 |
| Nicotinic acid | N0761-100G | HMDB0001488 | 123.0320 | 122.0242 | 124.0398 |
| O-Phosphoethanolamine | P0503-1G | HMDB0000224 | 141.0191 | 140.0113 | 142.0269 |
| Ornithine | O2375-5G | HMDB0000214 | 132.0899 | 131.0821 | 133.0977 |
| Orotic acid | O8402-25G | HMDB0000226 | 156.0171 | 155.0093 | 157.0249 |
| Oxalacetic acid | O4126-25G | HMDB0000223 | 132.0059 | 130.9981 | 133.0137 |
| Oxalic acid | 194131-5G | HMDB0002329 | 89.9953 | 88.9875 | 91.0031 |
| Oxidized glutathione | G4376-1G | HMDB0003337 | 612.1520 | 611.1442 | 613.1598 |
| Oxoglutaric acid | 75892-25G | HMDB0000208 | 146.0215 | 145.0137 | 147.0293 |
| Pantothenic acid | 21210-5G-F | HMDB0000210 | 219.1107 | 218.1029 | 220.1185 |
| Phenylpyruvic acid | P8001-5G | HMDB0000205 | 164.0473 | 163.0395 | 165.0551 |
| Phosphoenolpyruvic acid | P7127-250MG | HMDB0000263 | 167.9824 | 166.9746 | 168.9902 |
| Picolinic acid | P42800-100G | HMDB0002243 | 123.0320 | 122.0242 | 124.0398 |
| Putrescine | P7505-25G | HMDB0001414 | 88.1000 | 87.0922 | 89.1078 |
| Pyridoxal | P9130-5G | HMDB0001545 | 167.0582 | 166.0504 | 168.0660 |
| Pyridoxal 5'-phosphate | A3831.0005 | HMDB0001491 | 247.0246 | 246.0168 | 248.0324 |
| Pyridoxamine | P9380-1G | HMDB0001431 | 168.0899 | 167.0821 | 169.0977 |
| Pyridoxine | P9755-25G | HMDB0000239 | 169.0739 | 168.0661 | 170.0817 |
| Pyroglutamic acid | 83160-25G | HMDB0000267 | 129.0426 | 128.0348 | 130.0504 |
| Pyrophosphate | P8010 | HMDB0000250 | 177.9432 | 176.9354 | 178.9510 |
| Pyruvic acid | P8574-100G | HMDB0000243 | 88.0160 | 87.0082 | 89.0238 |
| Quinaldic acid | 160660-2.5G | HMDB0000842 | 173.0477 | 172.0399 | 174.0555 |
| Quinolinic acid | P63204-100G | HMDB0000232 | 167.0219 | 166.0141 | 168.0297 |
| Riboflavin | R4500-25G | HMDB0000244 | 376.1383 | 375.1305 | 377.1461 |
| Ribonic acid | 75406-10MG | HMDB0000867 | 166.0477 | 165.0399 | 167.0555 |
| Sarcosine | S7672-25G | HMDB0000271 | 89.0477 | 88.0399 | 90.0555 |
| Serotonin | H9523-25MG | HMDB0000259 | 176.0950 | 175.0872 | 177.1028 |
| Spermine | S1141-1G | HMDB0001256 | 202.2157 | 201.2079 | 203.2235 |
| Succinic acid | S3674-100G | HMDB0000254 | 118.0266 | 117.0188 | 119.0344 |
| Taurine | T0625-100G | HMDB0000251 | 125.0147 | 124.0069 | 126.0225 |
| Thiamine | T4625-25G | HMDB0000235 | 265.1123 | 264.1045 | 266.1201 |
| Thiamine monophosphate | T8637-5G | HMDB0002666 | 345.0786 | 344.0708 | 346.0864 |
| Thiamine pyrophosphate | C8754-5G | HMDB0001372 | 425.0450 | 424.0372 | 426.0528 |
| Thiosulfate | 72049-250G | HMDB0000257 | 113.9445 | 112.9367 | 114.9523 |
| Threonic acid | 380644-5G | HMDB0000943 | 136.0372 | 135.0294 | 137.0450 |
| Thymidine 5'-triphosphate | T0251-10MG | HMDB0001342 | 481.9893 | 480.9815 | 482.9971 |
| Thymine | T0376-5G | HMDB0000262 | 126.0429 | 125.0351 | 127.0507 |
| Tryptamine | 76706-100MG | HMDB0000303 | 160.1000 | 159.0922 | 161.1078 |
| Tryptophanol | T90301-5G | HMDB0003447 | 161.0841 | 160.0763 | 162.0919 |
| Uracil | U0750-5G | HMDB0000300 | 112.0273 | 111.0195 | 113.0351 |
| Uric acid | U2625 | HMDB0000289 | 168.0283 | 167.0205 | 169.0361 |
| Uridine | U3750-1G | HMDB0000296 | 244.0695 | 243.0617 | 245.0773 |

| HMDB Name | Reference | HMDB ID | Monoisotopic mass | $[M-H]^-$ | $[M+H]^+$ |
|---|---|---|---|---|---|
| Uridine 5'-diphosphate | 94330-100MG | HMDB0000295 | 404.0022 | 402.9944 | 405.0100 |
| Uridine 5'-monophosphate | U6375-1G | HMDB0000288 | 324.0359 | 323.0281 | 325.0437 |
| Uridine diphosphategalactose | U4500-10MG | HMDB0000302 | 566.0550 | 565.0472 | 567.0628 |
| Uridine triphosphate | U6750-100MG | HMDB0000285 | 483.9685 | 482.9607 | 484.9763 |
| Xanthurenic acid | D120804-5G | HMDB0000881 | 205.0375 | 204.0297 | 206.0453 |

***Table S2**. Criteria for assigning confidence in metabolite identification and corresponding terminology in the text. Table adapted from (Talavera Andujar et al. 2022)*

| Level | MS-DIAL - Non-target screening | Identification Term |
|---|---|---|
| **Level 1** | Confirmed structure by reference standard. MS, $MS^2$ and retention time match | Identified |
| **Level 2a** | ≥ 3 ion fragments matching; Dot product 70–100; Fragment presence 50–100 | Tentatively identified |
| **Level 3a** | ≥ 3 ion fragments matching; Dot product 50–70; Fragment presence 50–100 | Tentatively identified |

***Table S3**. List of polar metabolites, charge, retention time (RT), and confidence level for metabolite identification according to Table S2 and Talavera et al. (Talavera Andujar et al. 2022).*

| Metabolite | Charge | RT (min) | Identification level |
|---|---|---|---|
| Threonine | + | 9.87 | 1 |
| Serine | + | 10.86 | 1 |
| Glutamic acid | + | 11.57 | 1 |
| Glutathione | + | 11.33 | 1 |
| Glutamine | + | 10.30 | 1 |
| Tryptophan | + | 6.56 | 1 |
| Creatine | + | 9.51 | 1 |
| Phenylalanine | + | 4.95 | 1 |
| Acetylcarnitine | + | 4.63 | 1 |
| Aspartic acid | + | 11.90 | 1 |
| Choline | + | 4.31 | 1 |
| Proline | + | 7.15 | 1 |
| Methionine | + | 6.59 | 1 |
| Arginine | + | 13.18 | 1 |
| Cytosine | + | 4.52 | 1 |
| Adenine | + | 3.41 | 1 |
| Guanine | + | 6.42 | 1 |
| Inosine | + | 5.25 | 1 |
| Cytidine monophosphate | + | 12.39 | 1 |
| Adenosine 3'-monophosphate | + | 11.24 | 1 |
| Guanosine monophospate | + | 12.85 | 1 |
| Pyroglutamic acid | + | 11.57 | 1 |
| Niacinamide | + | 2.43 | 1 |

| | | | |
|---|---|---|---|
| Lactate | - | 7.11 | 1 |
| Tyrosine | - | 8.96 | 1 |
| Malic acid | - | 13.07 | 1 |
| Succinic acid | - | 12.06 | 1 |
| Taurine | - | 10.46 | 1 |
| Uridine 5'-monophosphate | - | 11.95 | 1 |
| Gamma-amino-n-butyric acid | + | 10.05 | 2a |
| Propionylcarnitine | + | 3.60 | 2a |
| Adenosine | + | 3.40 | 2a |
| Carnitine | + | 8.01 | 3a |
| Butyril carnitine | + | 2.99 | 3a |
| Citrulline | + | 10.78 | 3a |
| Guanosine | + | 7.59 | 3a |
| 3-Hydroxy-3-methylglutarate | - | 11.70 | 3a |
| Aconitic acid | - | 13.63 | 3a |
| Sorbitol | - | 8.31 | 3a |
| Acetylneuraminic acid | - | 10.65 | 3a |

***Table S4**. List of non-polar metabolites, charge, retention time (RT), and confidence level for metabolite identification according to Table S2 and Talavera et al. (Talavera Andujar et al. 2022).*

| Metabolite | Category | Charge | RT (min) | Identification level |
|---|---|---|---|---|
| PC(14:0/0:0) | Glycerophospholipids | + | 1.004 | 3 |
| PE(18:1(9Z)/0:0) | Glycerophospholipids | + | 1.871 | 3 |
| PE(18:0/0:0) | Glycerophospholipids | + | 2.736 | 3 |
| PC(16:1(9E)/0:0) | Glycerophospholipids | + | 1.106 | 3 |
| PC(0:0/16:0) | Glycerophospholipids | + | 1.634 | 3 |
| PC(18:1(9E)/0:0) | Glycerophospholipids | + | 1.664 | 3 |
| PC(20:4(8Z,11Z,14Z,17Z)/0:0) | Glycerophospholipids | + | 1.181 | 3 |
| PC(O-16:1(9Z)/20:4(8Z,11Z,14Z,17Z)) | Glycerophospholipids | + | 6.298 | 3 |
| Cer(d18:1/16:0) | Sphingolipids | + | 6.719 | 3 |
| Cer(d18:1/18:0) | Sphingolipids | + | 7.411 | 3 |
| Cer(d18:1/20:0) | Sphingolipids | + | 8.097 | 3 |
| Cer(d18:1/22:0) | Sphingolipids | + | 8.746 | 3 |
| Cer(d18:1(4E)/22:0(2OH)) | Sphingolipids | + | 8.088 | 3 |
| Cer(d18:1/24:1(15Z)) | Sphingolipids | + | 8.712 | 3 |
| Cer(d18:0/24:1(15Z)) | Sphingolipids | + | 8.941 | 3 |
| TG(16:0/16:0/16:0) | Glycerolipids | + | 11.841 | 3 |
| TG(16:0/18:1(9Z)/18:1(9Z))[iso3] | Glycerolipids | + | 11.861 | 3 |
| Coenzyme Q10 | Prenol Lipids | + | 10.428 | 3 |
| LPE O-16:1 | Lysophospholipid | - | 1.989 | 3 |
| Cholesterol sulfate | Sterol Lipids | - | 3.971 | 3 |
| Cer(d16:1/18:0) | Sphingolipids | - | 6.773 | 3 |
| Cer(d18:0/18:1) | Sphingolipids | - | 7.48 | 3 |
| SM 12:1;2O/20:0 | Sphingolipids | - | 5.228 | 3 |
| PE(18:1(9E)/18:1(9E)) | Glycerophospholipids | - | 6.923 | 3 |

| | | | | |
|---|---|---|---|---|
| PC(14:0/16:1(9Z)) | Glycerophospholipids | - | 5.487 | 3 |
| PE(P-18:0/22:6(4Z,7Z,10Z,13Z,16Z,19Z)) | Glycerophospholipids | - | 7.021 | 3 |
| PC(O-18:0/16:0) | Glycerophospholipids | - | 7.7 | 3 |
| PC(16:0/22:6(4E,7E,10E,13E,16E,19E) | Glycerophospholipids | - | 5.911 | 3 |
| PC(18:0/20:1(11Z)) | Glycerophospholipids | - | 7.96 | 3 |
| PC(18:0/22:6(4Z,7Z,10Z,13Z,16Z,19Z)) | Glycerophospholipids | - | 6.543 | 3 |
| PE(13:0/16:0) | Glycerophospholipids | - | 6.788 | 3 |
| PE(18:1(9Z)/0:0) | Glycerophospholipids | - | 1.9 | 3 |
| PE(20:4(8Z,11Z,14Z,17Z)/0:0) | Glycerophospholipids | - | 1.225 | 3 |
| PC(18:0/0:0) | Glycerophospholipids | - | 2.718 | 3 |
| PE(O-18:1(9Z)/0:0) | Glycerophospholipids | - | 3.189 | 3 |

**S2.1 Supplementary figures**

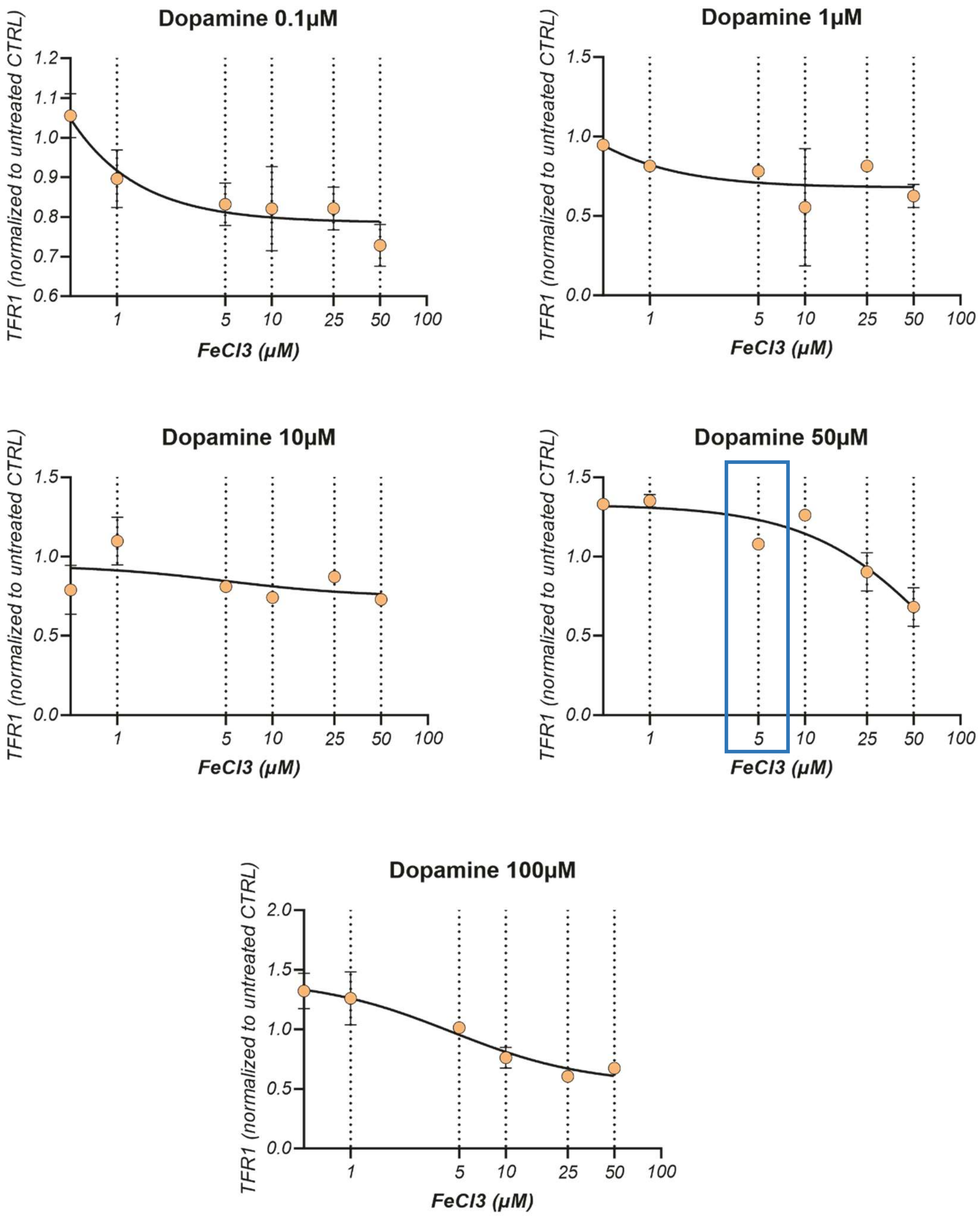

***Figure S1***. *Dose-response experiments for $FeCl_3$ and dopamine treatments in SH-SY5Y cells. Evaluation of TFR1 gene expression levels after 16 hours treatment with 0, 1, 5, 10, 25, 50, 100 µM $FeCl_3$ and 0.1, 10, 50, 100µM dopamine. 5 µM $FeCl_3$ and 50 µM dopamine were chosen for the experiments in this work.*

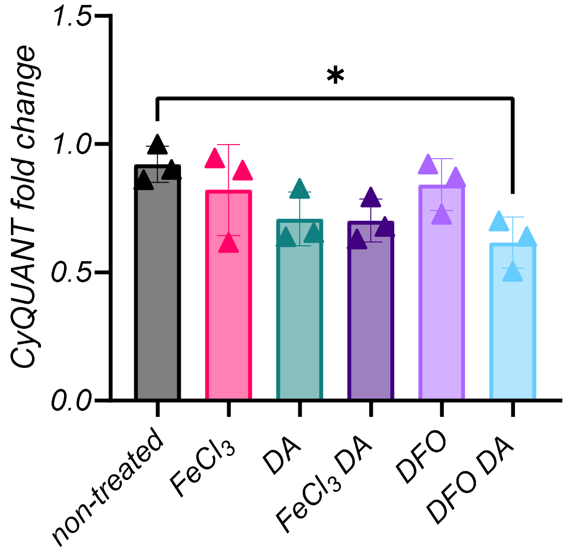

***Figure S2***. *Cell viability evaluated with the CyQUANT dye after 16 hours of treatments of SH-SY5Y cells. Effect on cell viability of dopamine (DA, 50 µM), iron ($FeCl_3$, 5 µM), deferoxamine (DFO, 5µM) and their respective combinations were evaluated. Data were normalized to non-treated condition; error bars represent SD of 3 data points; p-values were determined using one-way ANOVA followed by Dunnett's test to correct for multiple testing. *p≤0.05*

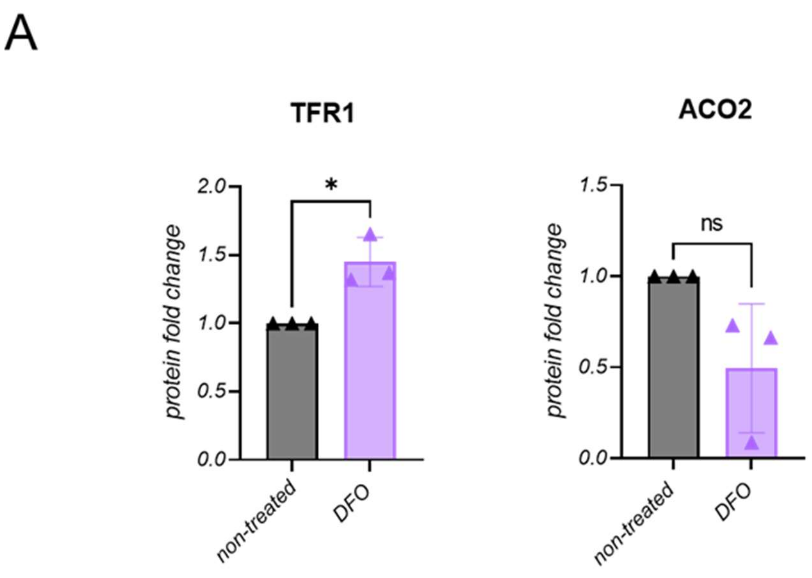

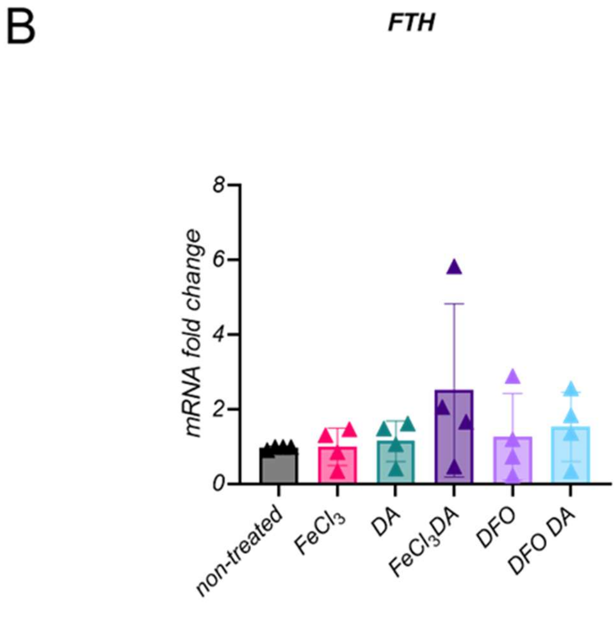

***Figure S3***. *Gene and protein expressions in SH-SY5Y cells. A) Densitometric analyses of TFR1 and ACO2 protein levels after treatment with DFO; n=3. B) mRNA expression levels of FTH after treatments; n=4. Statistical differences were calculated by Mann Whitney test. *p≤0.05.*

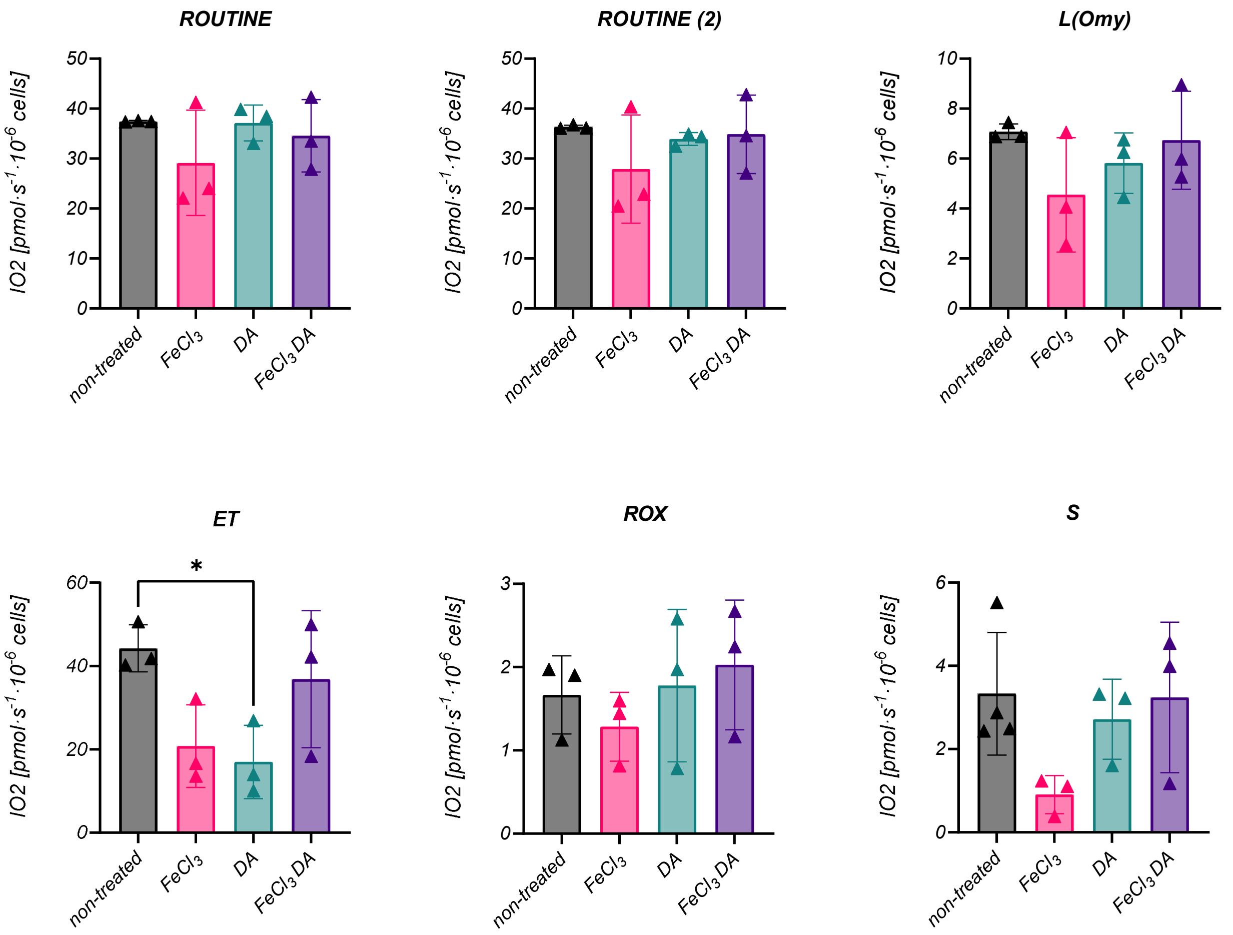

***Figure S4***. *Cell respiration measured at 16 hours of treatment in living SH-SY5Y neuroblastoma cells. Respiration values were normalized for the total number of cells per chamber. DA: dopamine. ROUTINE: ROUTINE respiration; ROUTINE (2): control of membrane integrity and permeability by pyruvate addition; L(Omy): non-phosphorylating resting state (LEAK state) after blocking the ATP synthase with oligomycin; ET: Electron transfer (ET) capacity (non-coupled ET state) after stepwise uncoupler additions; Rox: residual oxygen consumption in the ROX state due to oxidative side reactions; S: succinate used for viability test after inhibition of complex I with rotenone. Statistical analysis was carried out by using the ordinary one-way ANOVA followed by Dunnett's test to correct for multiple testing. *p≤0.05.*

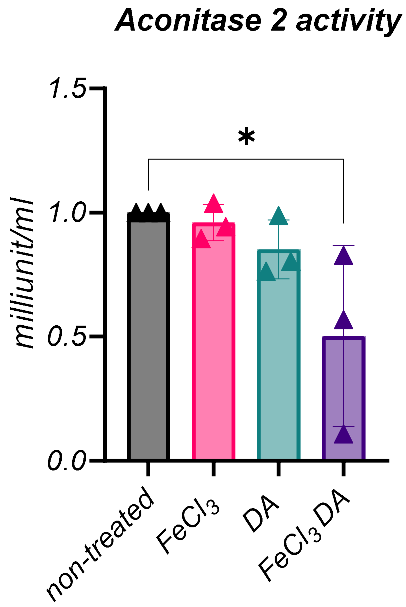

***Figure S5***. *Mitochondrial aconitase (ACO2) activity in SH-SY5Y cells upon treatments. DA: dopamine. Each condition was measured in n=3 independent assays. Statistical differences were calculated by ordinary one-way ANOVA followed by Dunnett's test to correct for multiple testing. *p≤0.05.*

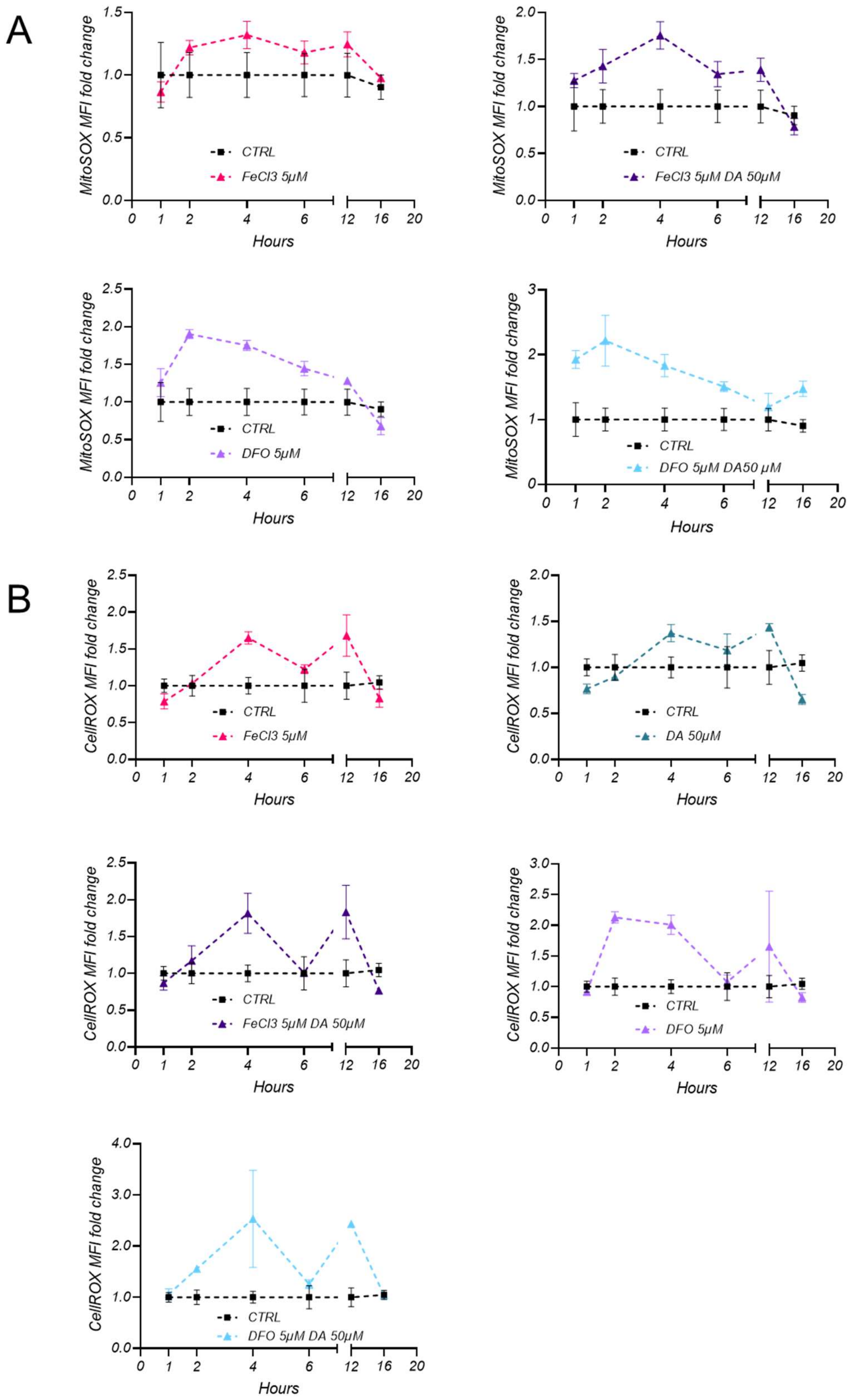

***Figure S6****. Fluctuation of ROS levels over time upon treatment of SH-SY5Y cells. A) Mitosox Red levels were quantified by FACS analysis after 1, 2, 4, 6, 12, and 16 hours. Cells treated with the different compounds are compared to cells without treatment as control. DA: dopamine. B) CellROX levels quantified by FACS after 1, 2, 4, 6, 12, and 16 hours of treatment. Cells treated with the different compounds are compared to cells without any treatment as control. DA: dopamine.*

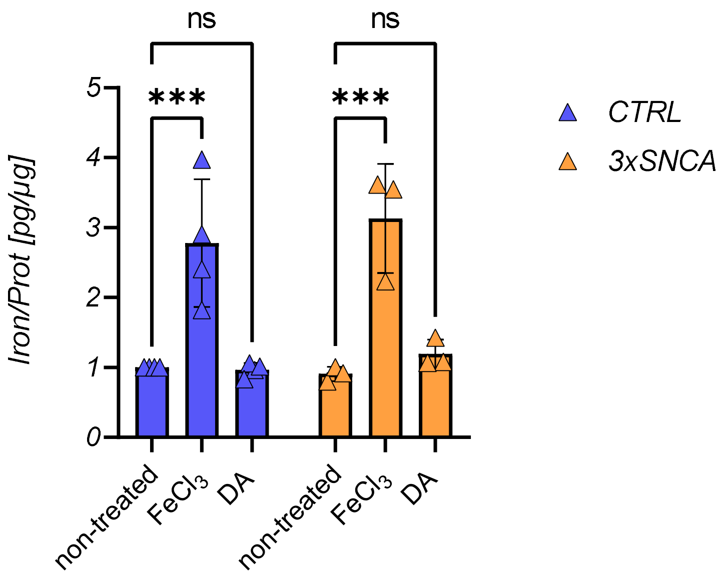

***Figure S7***. *Intracellular iron levels in iPSC-derived neurons of control and 3xSNCA patient lines. Cells were treated for 16 hours with iron or dopamine (DA) and intracellular iron levels were measured by atomic absorption spectroscopy. Results were normalized to protein content; n=3. Statistical differences were calculated by two-way ANOVA followed by Šídák's test to correct for multiple comparisons. ***p≤0.0001.*

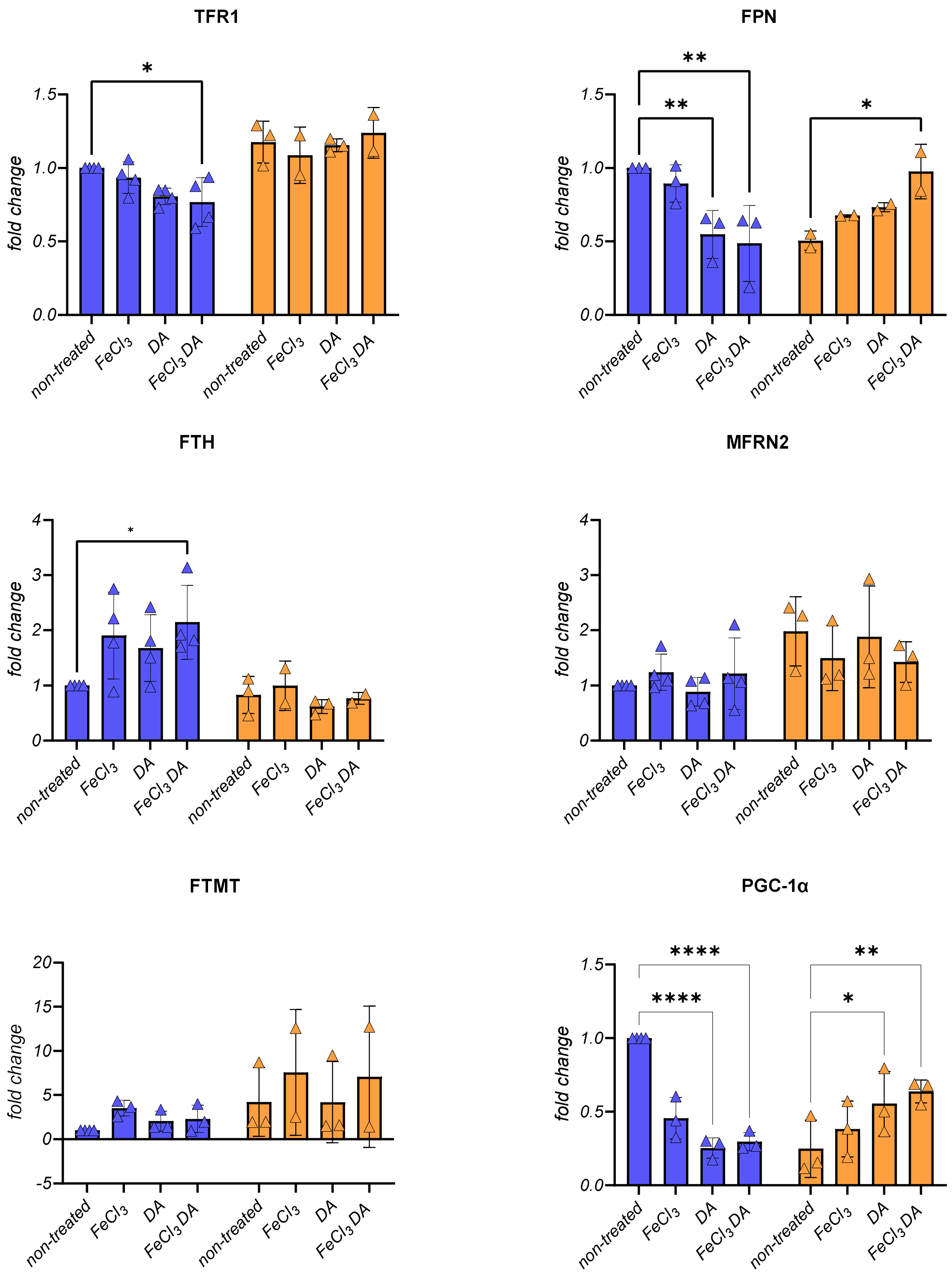

***Figure S8***. *Densitometric quantification of protein levels resolved by Western blot in iPSC-derived neurons. Levels of crucial proteins involved in iron import and storage in cells and mitochondria, in addition to markers of mitochondrial biogenesis. Two-way ANOVA was performed followed by Šídák's test to correct for multiple testing. *p≤0.05; **p≤0.01; ****p≤0.00001.*

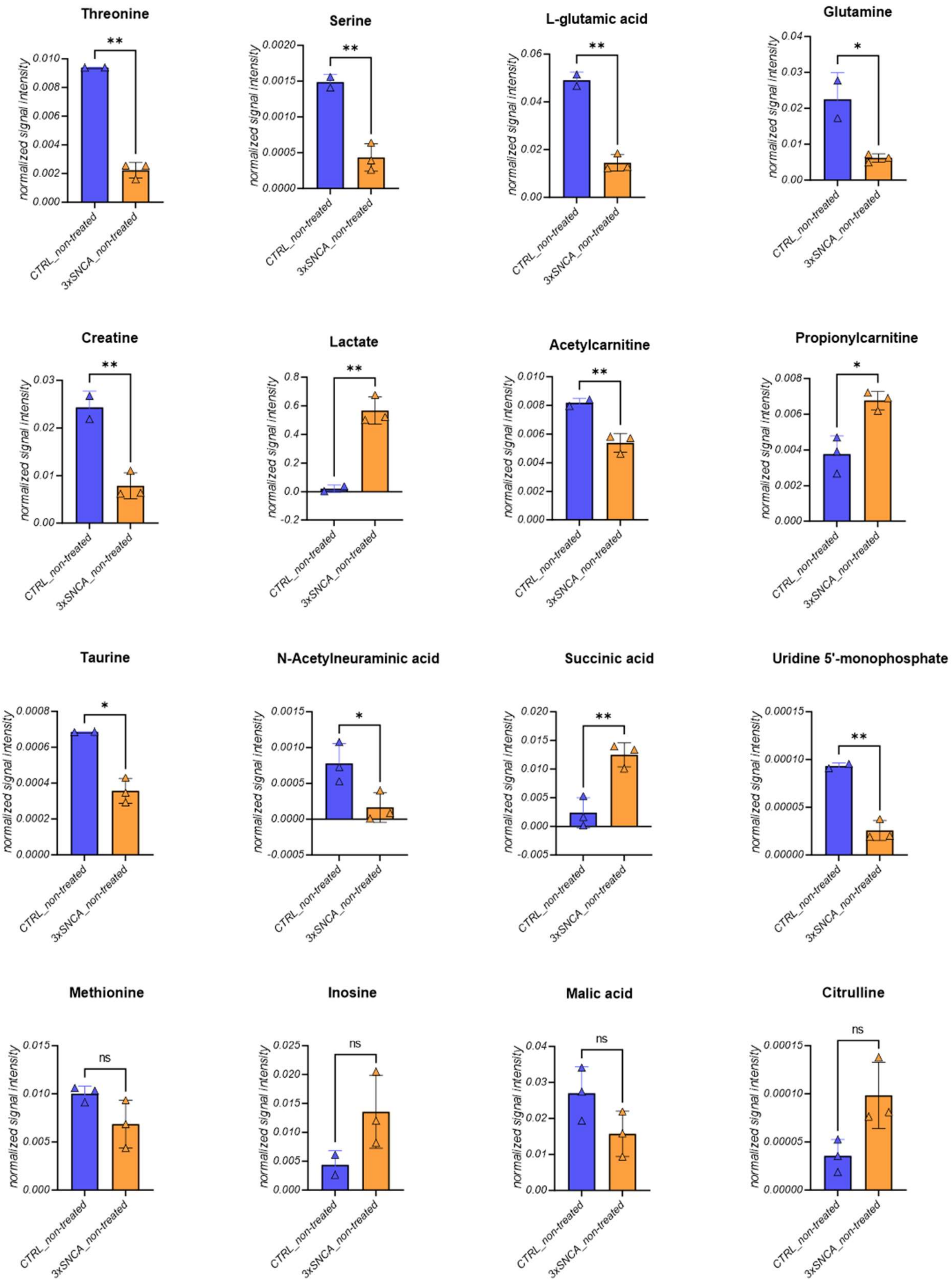

***Figure S9****. Quantification of polar metabolites in iPSC-derived control and patient neurons. Polar metabolites found to be present in significantly different amounts in the patient (3xSNCA) vs. control (CTRL) neurons or showing a trend towards increase or decrease in the PD patient line compared to the CTRL line. Statistical differences were calculated by unpaired t-test with Welch's correction. n≥2. *p≤0.05; **p≤0.01.*

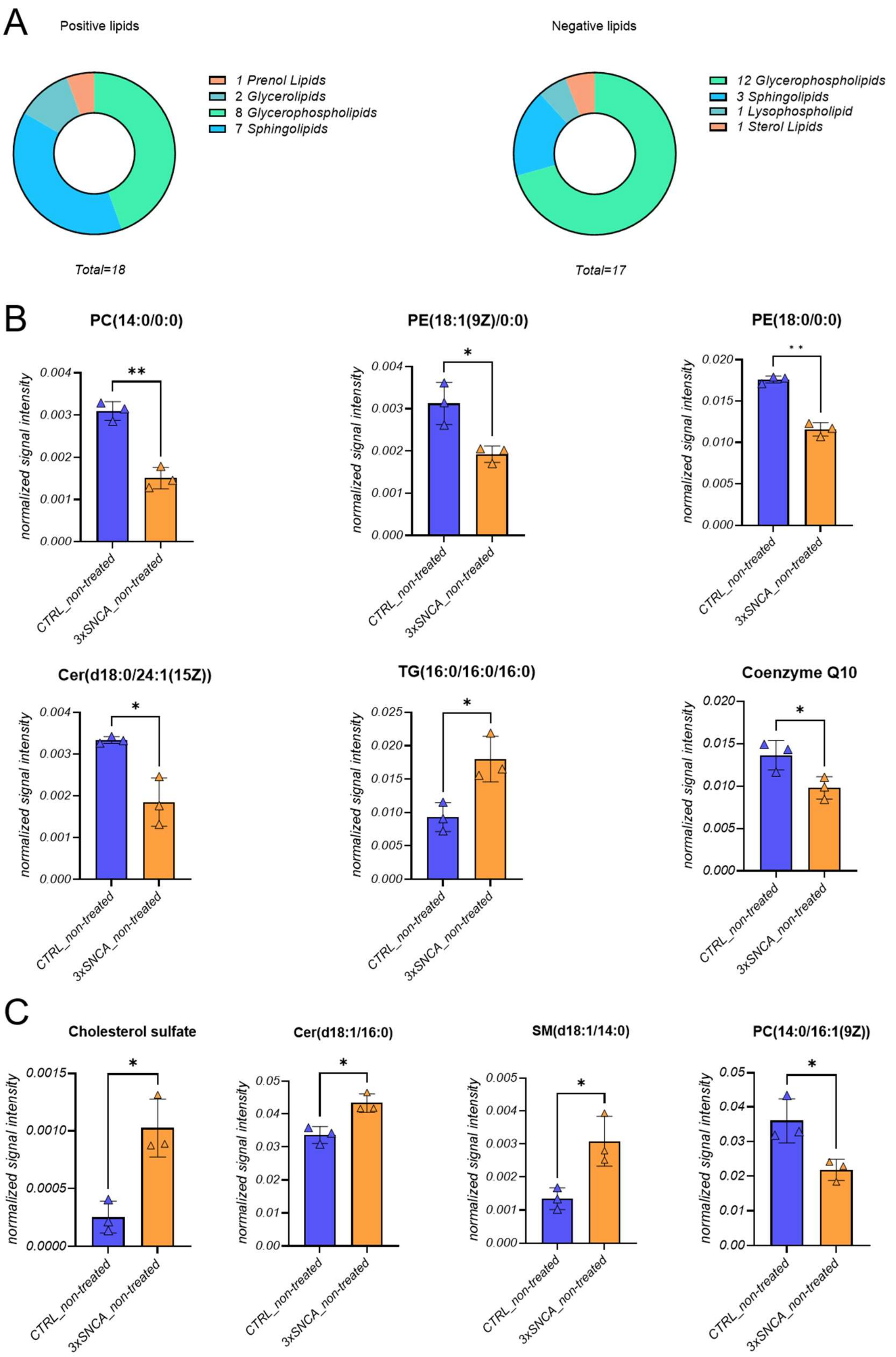

***Figure S10.*** *Non-polar metabolites detected in iPSC-derived control and patient neurons. A) Identified metabolites classified in classes and mode. B) Positively charged lipids differing between the two cell lines (CTRL vs 3xSNCA). C) Negatively charged lipids differing between the two cell lines (CTRL vs 3xSNCA). Statistical differences were calculated by unpaired t-test with Welch's correction. *p≤0.05; **p≤0.01.*

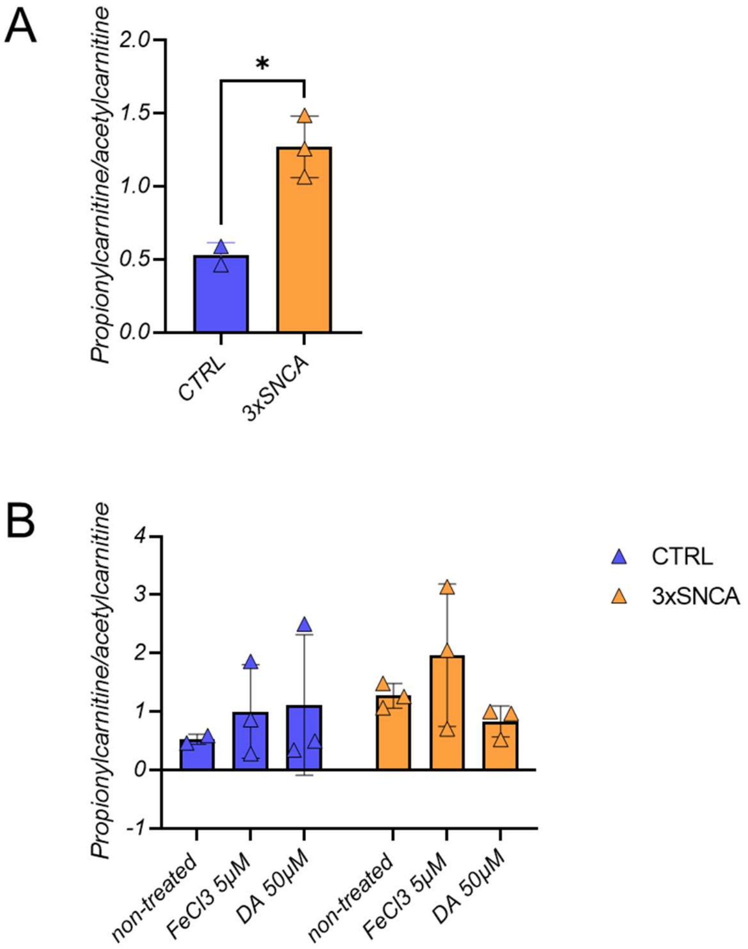

***Figure S11***. *Propionylcarnitine and acetylcarnitine ratio in iPSC-derived neurons. A) Comparison of propionylcarnitine/acetylcarnitine ratio between iPSC-derived neurons of CTRL and 3xSNCA lines. B) Alterations in propionylcarnitine/acetylcarnitine ratio induced by treatments in the two lines. *p≤0.05*

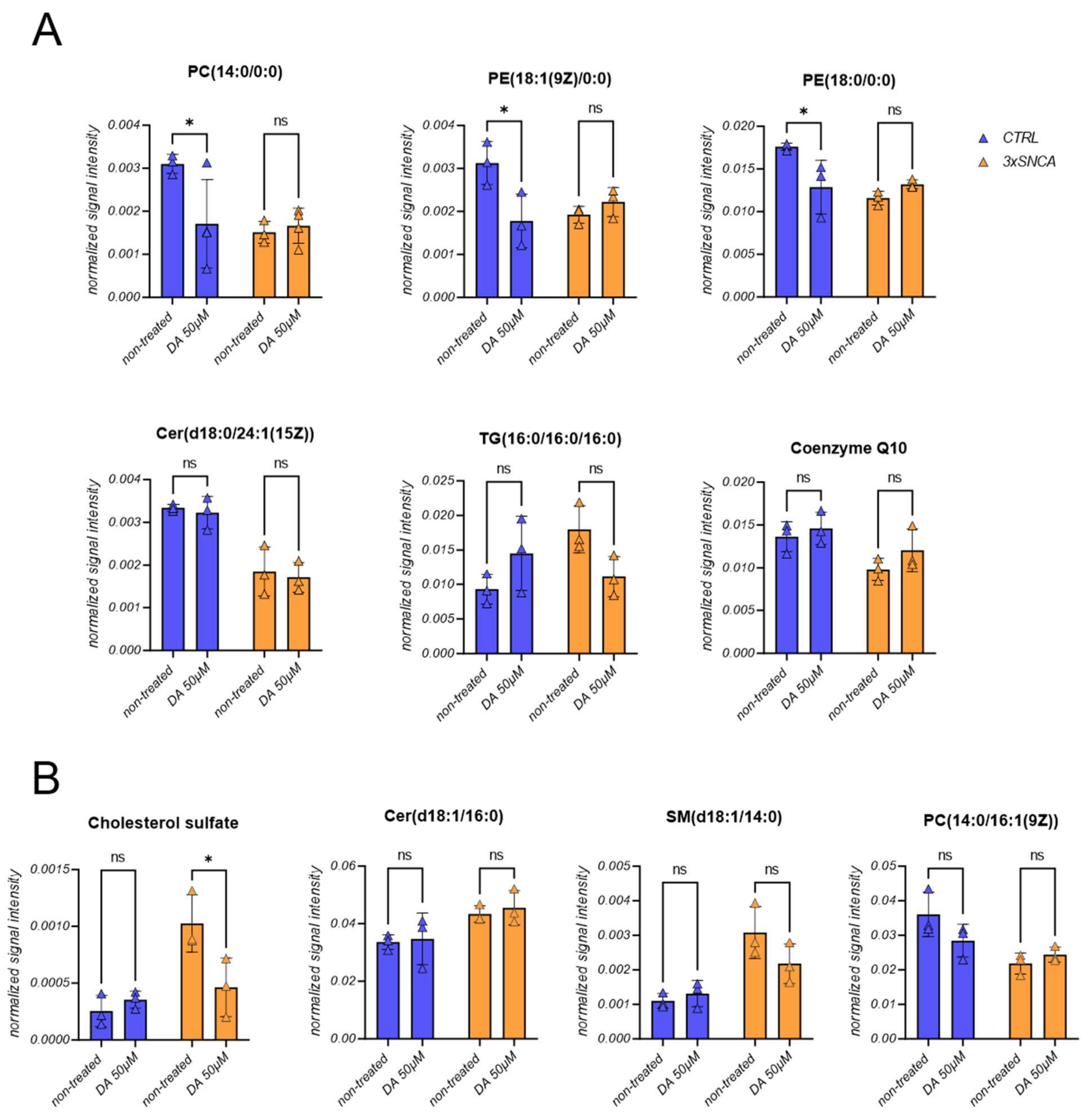

***Figure S12***. *Lipid levels upon dopamine treatment in iPSC-derived control and patient neurons. A) Positively charged lipids upon dopamine treatment. B) Negatively charged lipids upon dopamine treatment. Two-way ANOVA was performed followed by Šídák's test to correct for multiple testing. *p≤0.05.*

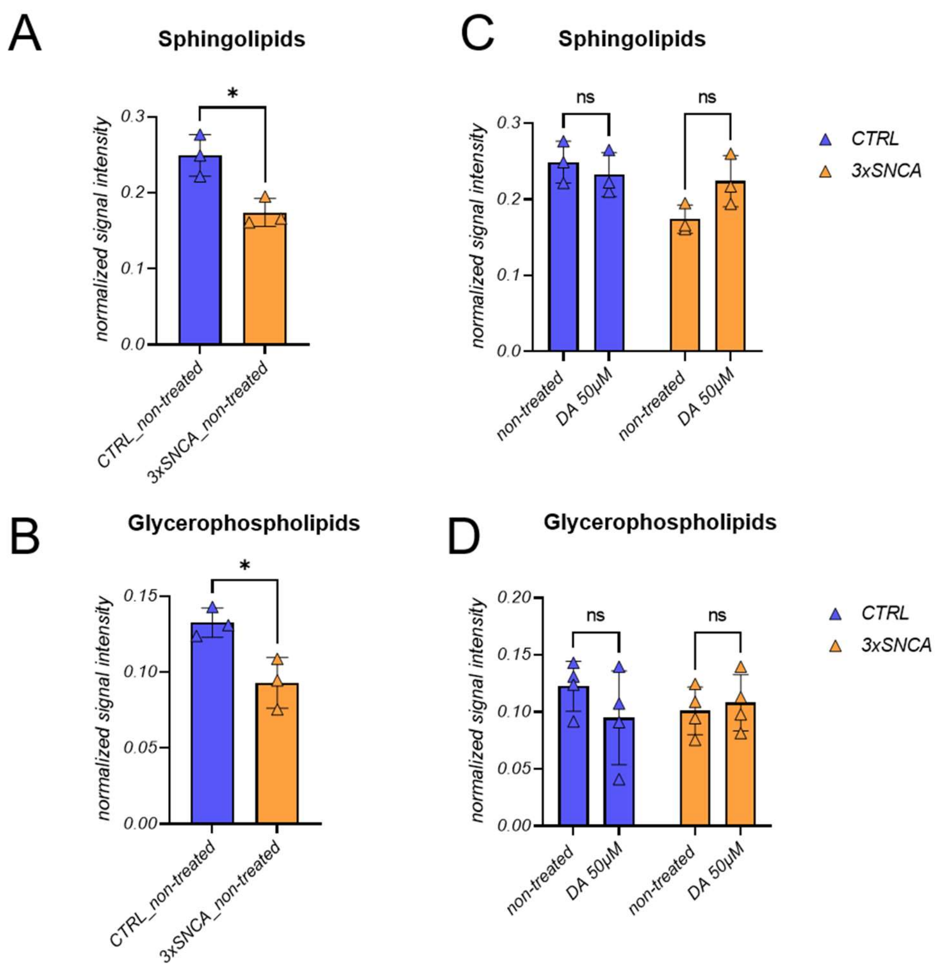

***Figure S13**. Analyses of lipid classes in iPSC-derived neuronal cell lines. A) and B) Classes of lipids changing between the control and PD patient-derived cell lines; C) and D) Levels of sphingolipids and glycerophospholipids after dopamine treatment in the two lines. Statistical differences were calculated by unpaired t-test with Welch's correction for A) and B) and two-way ANOVA followed by Šídák's test to correct for multiple testing for C) and D). *p≤0.05.*

#### *S3 References*

Cajka, T., J. T. Smilowitz, and O. Fiehn. 2017. 'Validating Quantitative Untargeted Lipidomics Across Nine Liquid Chromatography-High-Resolution Mass Spectrometry Platforms', *Anal Chem*, 89: 12360-68.

Talavera Andujar, B., D. Aurich, V. T. E. Aho, R. R. Singh, T. Cheng, L. Zaslavsky, E. E. Bolton, B. Mollenhauer, P. Wilmes, and E. L. Schymanski. 2022. 'Studying the Parkinson's disease metabolome and exposome in biological samples through different analytical and cheminformatics approaches: a pilot study', *Anal Bioanal Chem*, 414: 7399-419.
